# The polyglutamine amyloid nucleus in living cells is a monomer with competing dimensions of order

**DOI:** 10.1101/2021.08.29.458132

**Authors:** Tej Kandola, Shriram Venkatesan, Jiahui Zhang, Brooklyn Lerbakken, Jillian F Blanck, Jianzheng Wu, Jay Unruh, Paula Berry, Jeffrey J. Lange, Alex Von Schulze, Andrew Box, Malcolm Cook, Celeste Sagui, Randal Halfmann

**Author notes:** These authors contributed equally (Tej Kandola, Shriram Venkatesan).

## Abstract

A long-standing goal of the study of amyloids has been to characterize the structural basis of the rate-determining nucleating event. However, the ephemeral nature of that event has made it inaccessible to classical biochemistry, structural biology, and computational approaches. Here, we addressed that limitation by measuring the dependence of amyloid formation on concentration and conformational templates in living cells, whose volumes are sufficiently small to resolve the outcomes of independent nucleation events. We characterized numerous rationally designed sequence variants of polyglutamine (polyQ), a polypeptide that precipitates Huntington’s and other amyloid-associated neurodegenerative diseases when its length exceeds a characteristic threshold. This effort uncovered a pattern of approximately twelve Qs, only for polypeptides exceeding the clinical length threshold, that allow for amyloid nucleation to occur spontaneously within single polypeptides. Nucleation was inhibited by intermolecular phase separation. Using atomistic molecular dynamics simulations, we found that the pattern encodes a minimal steric zipper of interdigitated side chains. Lateral growth of the steric zipper competed with axial growth to produce “pre-amyloid” oligomers. By illuminating the structural mechanism of polyQ amyloid formation in cells, our findings reveal a potential molecular etiology for polyQ diseases, and may provide a roadmap for the design of new therapies.

## Introduction

Amyloids are highly ordered protein aggregates. They are widely associated with aging and neurodegenerative diseases (Chiti and Dobson, 2017; Huang et al., 2019), but also have functional roles in programmed cell death and subcellular compartmentalization (Boke et al., 2016; Hervas et al., 2020; Maji et al., 2009; Majumdar et al., 2012; O’Carroll et al., 2020; Rodríguez Gama et al., 2021; Vogler et al., 2018). These activities derive from an inherent tendency of amyloids to catalyze their own formation, a phenomenon driven by supersaturation resulting from sequence-encoded kinetic barriers to nucleation. The structural basis of nucleation remains mysterious despite decades of intense research.

Polyglutamine (polyQ) is an amyloid-forming module common to eukaryotic proteomes (Mier et al., 2020). In humans, it is responsible for nine invariably fatal neurodegenerative diseases, the most prevalent of which is Huntington’s Disease. A remarkable feature of polyQ pathology -- found equally in Huntington’s disease patients, cell culture, organotypic brain slices, and animal models -- is that neuronal loss occurs with a constant frequency over time, implying that death happens independently and stochastically in each cell (Clarke et al., 2000; Linsley et al., 2019). Two consistent observations suggest that this feature emerges directly from the nucleation barrier. First, the kinetics of polyQ aggregation in vitro, in cells, and in organismal models is consistent with primary nucleation as the rate-limiting step (Chen et al., 2002; Colby et al., 2006; Kakkar et al., 2016; Sinnige et al., 2021). Second, polyQ diseases result almost exclusively from an *intramolecular* change, specifically, an expansion in the number of sequential glutamines beyond a protein-specific threshold centered around 38 residues (Lieberman et al., 2019). In contrast to other amyloid diseases, which are frequently brought on by gene duplication, misplicing, and post-translational processing (Book et al., 2018; Kim et al., 2020; Selkoe and Hardy, 2016), polyQ diseases have relatively few *intermolecular* modifiers (Lee et al., 2019; Wexler et al., 1987). Hence, more so than for any other proteopathy, progress against polyQ diseases awaits detailed knowledge of the amyloid nucleus itself.

Despite its extraordinary relevance to both basic biology and human health, no empirically determined structure of an amyloid nucleus yet exists. This is because nuclei cannot be observed directly by any existing experimental approach. Unlike mature amyloid fibrils, which are stable and amenable to structural biology, nuclei are by definition unstable. Their structures do not necessarily persist into the mature amyloids that arise from them (Auer et al., 2008; Buell, 2017; Hsieh et al., 2017; Levin et al., 2014; Liang et al., 2018; Li et al., 2010; Sil et al., 2018; Yamaguchi et al., 2005; Zanjani et al., 2020). Nucleation occurs far too infrequently, and involves far too many degrees of freedom, to simulate computationally from a naive state (Barrera et al., 2021; Kar et al., 2011; Strodel, 2021). Unlike for phase separation, whose nucleation concerns primarily a loss of *intermolecular* entropy, amyloid nucleation also involves a major loss of *intramolecular* entropy (**Fig. S1A**), i.e. selects for a specific combination of backbone and side chain torsion angles (Khan et al., 2018; Vitalis and Pappu, 2011; Zhang and Schmit, 2016). As a consequence of the latter requirement, amyloid-forming proteins can accumulate to supersaturating concentrations while still remaining soluble, thereby storing potential energy that will subsequently drive their aggregation following a stochastic nucleation event (Buell, 2017; Khan et al., 2018). Once a nucleus does form, additional molecules join with relative ease, depleting the soluble level of the protein and changing its activity throughout the cell or organism. It is for this reason that amyloid phenomena tend to progress toward predetermined fates, such as programmed cell death (Majumdar et al., 2012; Rodríguez Gama et al., 2021), neurodegeneration and aging (Chiti and Dobson, 2017; Huang et al., 2019).

Due to the improbability of critical fluctuations occurring simultaneously in both density and conformation (i.e. *homogeneous* nucleation), amyloid nucleation tends to involve *heterogeneities,* or metastable intermediates of varying stoichiometry and conformation (Auer et al., 2008; Buell, 2017; Hsieh et al., 2017; Levin et al., 2014; Liang et al., 2018; Li et al., 2010; Serio et al., 2000; Sil et al., 2018; Vitalis and Pappu, 2011; Yamaguchi et al., 2005; Zanjani et al., 2020). These heterogeneities erode the barrier to nucleation. The occurrence of heterogeneities in the nucleation pathway also implies that the nature of the rate-limiting structure for a given amyloid will depend both on the protein’s *concentration* (**Fig. S1B**) and the existence of other cellular factors that influence the protein’s *conformation* (**Fig. S1C**) (Bradley et al., 2002; Buell, 2017; Collinge and Clarke, 2007; Sanders et al., 2014; Törnquist et al., 2018).

In principle, however, the sequence-encoded structural features of an amyloid nucleus may be deduced by studying the effects on nucleation of iterative rational mutations made to the nucleating polypeptide. Mutations that raise or lower the barrier independently of concentration provide information by which to discriminate the possible structural models. With enough constraints, the conformation representing the saddle point of the lowest energy path over the nucleation barrier, will emerge. In practice, this approach would require an assay capable of distinguishing kinetic from equilibrium effects on amyloid formation for a large number of mutations. Classic assays of *in vitro* amyloid assembly kinetics cannot achieve this goal, both because of their limited throughput, and because experimentally tractable reaction volumes are far too large for nucleation to be rate-limiting for biologically relevant amyloids (Michaels et al., 2017).

We recently developed an assay to circumvent these limitations (Khan et al., 2018; Posey et al., 2021; Venkatesan et al., 2019). Distributed Amphifluoric FRET (DAmFRET) uses a photoconvertible fusion tag and high throughput flow cytometry to treat living cells as femtoliter-volume test tubes, thereby providing the large numbers of independent reaction vessels of exceptionally small volume that are required to discriminate independent nucleation events under physiological conditions (**Fig. 1A**). Compared to conventional reaction volumes for protein self-assembly assays, the budding yeast cells employed in DAmFRET increase the dependence of amyloid formation on nucleation by nine orders of magnitude (the difference in volumes). DAmFRET employs a high-variance expression system to induce a hundred-fold range of concentrations of the protein of interest. By taking a single snapshot of the protein’s extent of self-association in each cell, at each concentration, at a single time point appropriate for the timescale of amyloid formation (hours), we probe the magnitudes of critical fluctuations in both density and conformation that govern nucleation.

**Figure 1.**
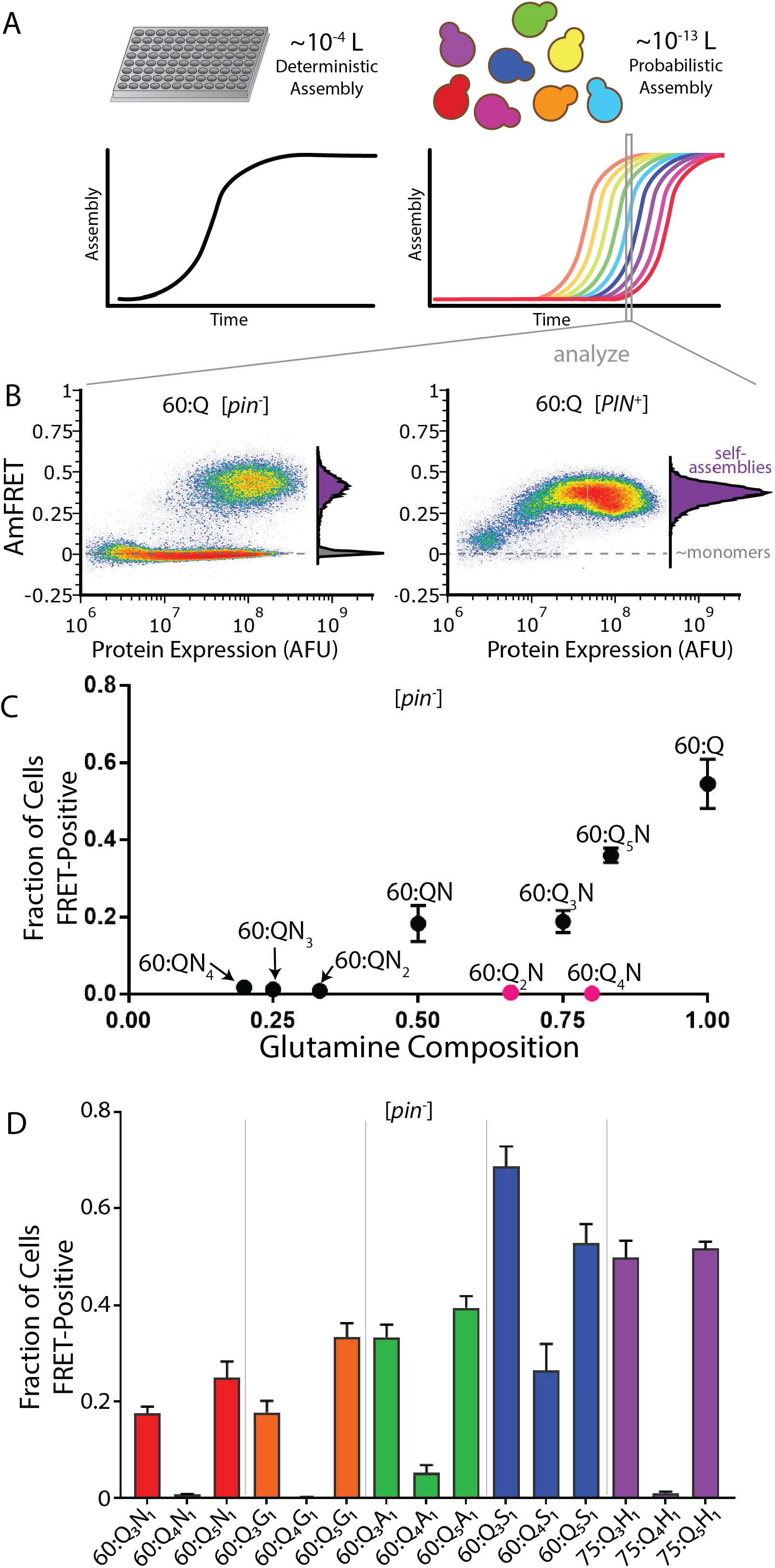
A cryptic pattern of glutamines governs amyloid nucleation. A. Volumes of reaction vessels influence the kinetics of amyloid formation. The concentrations of amyloid nuclei are so low that amyloid formation is rate-limited by nucleation in the femtoliter volumes of individual yeast cells (right) but not in the microliter volumes of conventional in vitro kinetic assays (left). B. DAmFRET provides a snapshot of the progress of amyloid formation as a function of expression level in each of several thousand cells at a single time point. DAmFRET profiles, showing the extent of self-assembly (AmFRET) as a function of protein expression level (AFU, arbitrary fluorescence units) of polyglutamine (60:Q) in the absence ([*pin^-^*]) and presence ([*PIN^+^*]) of endogenous amyloids of a different Q/N-rich protein. PolyQ has a large nucleation barrier in [*pin*^-^] cells, as evidenced by the window of concentration with a bimodal distribution of AmFRET values, wherein cells either lack (no AmFRET) or contain (high AmFRET) amyloid. [*PIN^+^*] eliminates the nucleation barrier, as evidenced by all cells shifting to the high AmFRET population. Insets show histograms of AmFRET values. C. Scatter plot of the fraction of [*pin*^-^] cells with amyloid versus the fraction of glutamines in the protein’s sequence. Red dots denote repeats with an even number of glutamines followed by an asparagine. Shown are means +/- SD of biological quadruplicates (independent yeast transformants). D. Bar plot of the fraction of [*pin*^-^] cells with amyloid for Q_3_X, Q_4_X, and Q_5_X sequences where X = N, G, A, S, or H. Shown are means +/- SD of biological quadruplicates. *Length of the QxH sequences is 75; all others are 60.

The yeast system additionally allows for orthogonal experimental control over the critical conformational fluctuation. This is because yeast cells normally contain exactly one cytosolic amyloid species -- a prion state formed by the low abundance Q- and N-rich endogenous protein, Rnq1 (Kryndushkin et al., 2013; Nizhnikov et al., 2014). The prion state can be gained or eliminated experimentally to produce cells whose sole difference is whether the Rnq1 protein does ([*PIN^+^*]) or does not ([*pin*^-^]) exist in an amyloid state (Derkatch et al., 2001). [*PIN^+^*] serves as an imperfect template that stabilizes amyloid conformations in compositionally similar proteins. Known as cross-seeding, this phenomenon is believed to be analogous to, but much less efficient than, homotypic amyloid elongation (Keefer et al., 2017; Khan et al., 2018; Serio, 2018). By evaluating nucleation frequencies as a function of concentration in both [*pin*^-^] and [*PIN*^+^] cells, we uncouple the two components of the nucleation barrier and thereby relate specific sequence features to the nucleating conformation.

Thus equipped, we sought to deduce the structure of the polyQ amyloid nucleus. Fortunately, polyQ may be an ideal model to elucidate nucleus structure. It has zero sequence complexity, which profoundly simplifies the design and interpretation of sequence variants. Additionally, unlike other pathogenic protein amyloids, which are notoriously polymorphic at a structural level, polyQ amyloids have an invariant core structure even under different assembly conditions and with different flanking domains (Boatz et al., 2020; Galaz-Montoya et al., 2021; Lin et al., 2017). This means that amyloid nucleation, and pathogenesis, of polyQ emerges directly from its ability to spontaneously acquire a *specific* nucleating conformation, and any variant that decelerates nucleation can be interpreted with respect to its effect on *that* conformation.

## Results

### A cryptic pattern of glutamines governs amyloid nucleation

We used our DAmFRET approach to quantify polyQ nucleation events among approximately one hundred thousand cells, per experiment, as a function of two variables that separately govern the density and conformational components of the nucleation barrier -- concentration and Rnq1 amyloid status.

In [*pin*^-^] cells, we found that cells expressing polyQ partitioned into two discontinuous populations: one containing only monomers, as indicated by an absence of AmFRET **(Fig. 1B)**, and the other containing self-assemblies of the protein, as indicated by high AmFRET **(Fig. 1B)**. The two populations occurred in overlapping windows of expression (~10^7^-10^8^ AFU, Fig. 2B left), with very few cells at intermediate AmFRET levels. This pronounced discontinuity in the relationship between self-assembly and concentration is characteristic for prion-like amyloids, and confirms the existence of a rate-limiting conformational fluctuation. Once nucleation occurs, however, the protein rapidly polymerizes to a steady state level (Buell, 2017; Khan et al., 2018).

**Figure 2.**
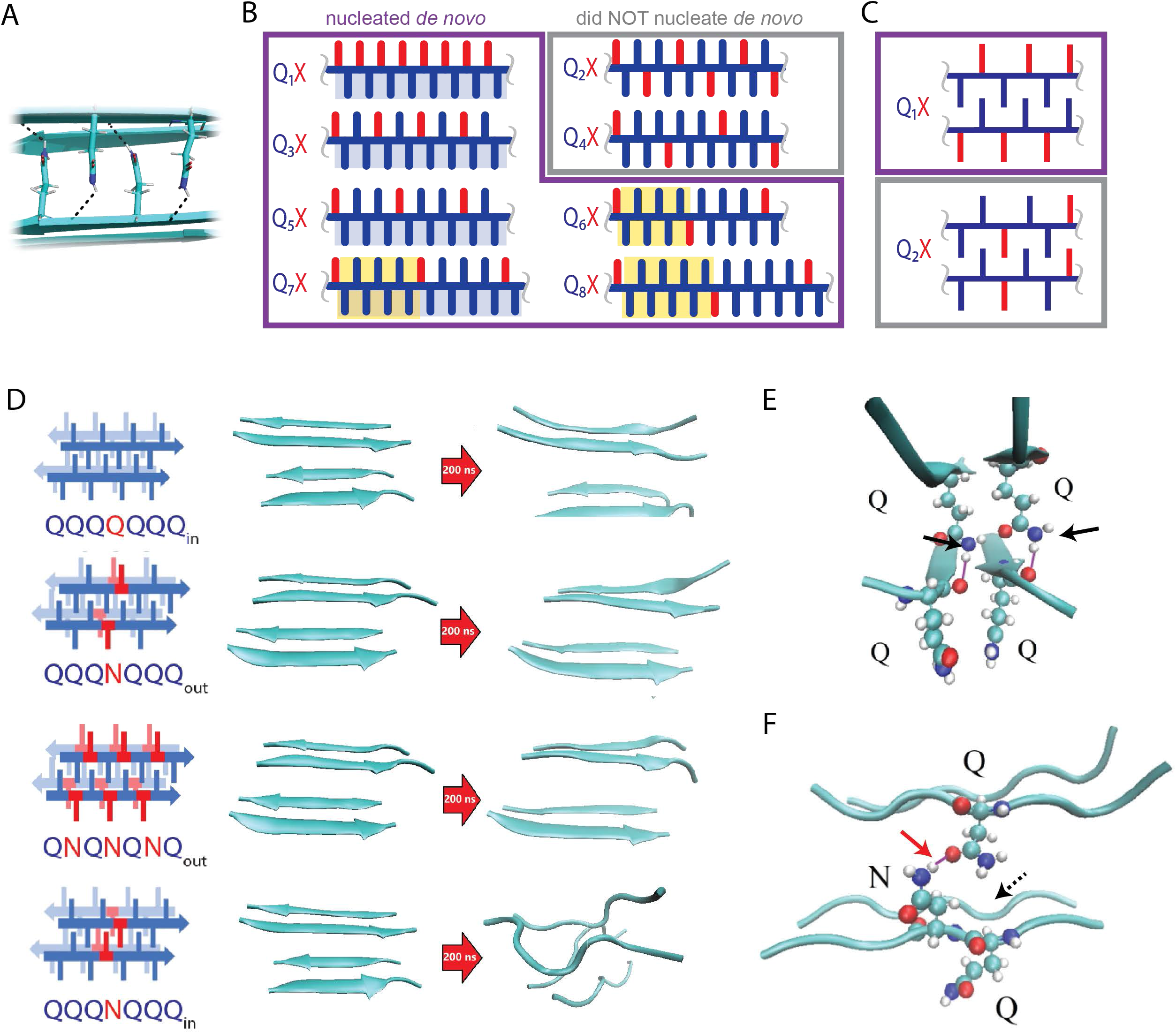
Unilaterally contiguous glutamines encode an exclusive steric zipper structure. A. A view down the axis of a local segment of all-glutamine steric zipper, or “Q zipper”, between two antiparallel two-stranded sheets. Only the top four internally facing side-chains are shown, to emphasize interdigitation and H-bonding between the terminal amides and the opposing backbone. B. Schema showing side chain arrangements along a continuous beta strand for sequences composed of tandem repeats of Qs (blue) interrupted by single non-Q residues (X, red). Shading indicates contiguous stretches of Qs on one (blue) or both (yellow) sides of the strand. Note that bilateral contiguity is bounded by X on either side of the strand. Note also that the illustrated strands will not necessarily be continuous in the context of the nucleus; i.e. parts of the sequence in the 60 residue-long polypeptide must form loops that interconnect strand elements. C. Schema of the tertiary contacts between two beta strands, as in a steric zipper. The zipper can be formed only from Q residues for repeats that have an odd number of Qs (e.g. Q_1_X), but not for repeats with an even number of Qs (e.g. Q_2_X). D. Molecular simulations of model Q zippers formed by a pair of two-stranded antiparallel beta sheets, wherein non-Q residues (in red) face either inward or outward. The schema are oriented so the viewer is looking down the axis between two sheets. The zipper is stable for pure polyQ (QQQQQQQ, top simulation), or when substitutions face outward (QQQNQQQ, second simulation; and QNQNQNQ, fourth simulation), but not when substitutions face inward (QQQNQQQ, third simulation). E. Snapshot from the uninterrupted Q zipper simulation, showing H-bonds (black arrows) between internal extended Q side chains and the opposing backbones. F. Snapshot from the internally interrupted Q zipper simulation, illustrating that the side chain of N is too short to H-bond the opposing backbone. However, the N side chain is long enough to H-bond the opposing Q side chain (red arrow), thereby intercepting the side chain-backbone H-bond that would otherwise occur (dashed arrow) between that Q side chain and the backbone amide adjacent to the N. This leads to dissolution of the zipper.

The tendency of Q/N-rich proteins to form amyloid increases with their length (Alberti et al., 2009). That polyQ nucleated robustly even at the relatively short length of 60 distinguishes it from dozens of physicochemically similar (Q-rich) amyloids now characterized by DAmFRET (Khan et al., 2018; Posey et al., 2021) and unpublished. This suggests that nucleation is promoted either by the exceptionally low sequence complexity of pure polyQ and/or by an emergent property of Q side chains, specifically. To discriminate between these possibilities, we tested an equivalent length homopolymer of asparagine, whose side chain is one methylene shorter but otherwise identical to that of glutamine. Remarkably, the protein failed to acquire AmFRET even at the highest level of expression (**Fig. S2A**, 60:N, [*pin*^-^]) -- approximately 200 μM (Khan et al., 2018). To determine if this can be attributed to a larger conformational nucleation barrier than that of polyQ, we expressed both proteins in cells harboring the heterogeneous conformational template, [*PIN^+^*], Indeed, polyN now nucleated with a comparable frequency and concentration-dependence to well-characterized Q/N-rich prion-forming proteins (compare **Fig. S2A**, 60:N, [*PIN^+^*] with plots in Posey et al. 2021). Interestingly, for polyQ, the overlapping expression range between monomer- and amyloid-containing cell populations was eliminated by [*PIN^+^*], indicating that polyQ is exceptionally susceptible to this entity (**Fig. 1B**, 60:Q, [*PIN^+^*]). We conclude that the polyQ amyloid nucleus contains a conformational element that is stabilized by cooperative interactions unique to Q side chains.

The surprising inability of N to form this nucleating element, despite its being the most physicochemically similar residue to Q, led us to consider whether a genetic screen of N-substituted polyQ variants might reveal patterns that uniquely encode the structure of the nucleus. We reasoned that a random screen of such sequences would be unlikely to yield informative patterns, however, given that the universe of Q/N sequences, even if limited to just 40 residues (slightly above the pathogenic length threshold for Huntington’s Disease), exceeds one trillion.

We therefore employed a systematic approach to rationally sample Q/N sequence space. Each sequence in the resulting series comprises a tandem repeat of *a* Qs separated by *b* Ns for a total length of *L* residues (L:Q_a_N_b_, where *a* and/or *b* = 1; **Table S1**). We characterized the self-assembly of each sequence via DAmFRET in both [*pin*^-^] and [*PIN*^+^] cells. The resulting dataset revealed three sequence determinants of amyloid nucleation.

First, repeats that were at least 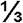 Ns (i.e. for *a* ≤ 2) resembled polyN rather than polyQ: they nucleated in a highly concentration-dependent fashion, and only in [*PIN*^+^] cells (**Fig. S2A**). This type of amyloid is clearly distinct from that of polyQ, perhaps reflecting the preference of polyN for parallel rather than antiparallel arrangements (Zhang et al., 2016). We did not characterize it further here. Second, de novo nucleation only occurred for repeats with at least 50% Qs (i.e. for *b* = 1). Third, and quite unexpectedly, [*pin*^-^] nucleation just above this compositional threshold only occured for the subset of repeats where *a* is an odd value (60:Q_<u>1</u>_N_1_, 60:Q_<u>3</u>_N_1_, 60:Q_<u>5</u>_N_1_), and not for those where *a* is an even value (60:Q_<u>2</u>_N_1_, 60:Q_<u>4</u>_N_1_) (**Fig. 1C** and see boxed region in **S2B**). In other words, while nucleation increases with Q density, there exists a rule related to the spacing of Qs in the sequence.

To determine if this rule concerns the glutamine side chains themselves, rather than their interaction with asparagines, we uniformly substituted Ns with alternative residues (i.e. L:Q_a_X_1_, with the subscript “1” omitted hereafter) of diverse physicochemistry -- either glycine, alanine, serine, or histidine. In all cases, the even/odd dependence persisted. That is, de novo nucleation was much more frequent for repeats of Q_3_X and Q_5_X (hereafter, Q_(3,5)_X) than for Q_4_X (**Fig. 1D, S2C**). The difference was, however, less pronounced in the case of S, all three repeat variants of which nucleated more robustly than for other identities of X. Unlike Q_4_N and Q_4_G, the Q_4_S and to a lesser extent Q_4_A and Q_4_H variants did detectably nucleate de novo, albeit in a concentration-dependent manner resembling that of longer even-numbered variants (Q_(6+)_N). The trends persisted, with Q_(3,5)_X nucleating better than Q_4_X, even in the presence of a heterogeneous conformational template, [*PIN^+^*] **(S2D).**

### Unilaterally contiguous glutamines encode an exclusive zipper structure

As a starting point to uncover possible physical explanations for the dependence on even/odd spacing, we analyzed representative sequences -- 60:Q_3_N, 60:Q_4_N, 60:Q_5_N, 60:Q, 60:N -- with state-of-the-art amyloid predictors (Charoenkwan et al., 2021; Keresztes et al., 2021; Prabakaran et al., 2021). Using their respective default parameters, we found that most predictors failed entirely to detect amyloid propensity among these sequences. Remarkably, none of the predictors distinguished 60:Q_4_N relative to 60:Q_(3,5)_N (Table S2), despite their completely dissimilar amyloid propensities. Apparently, the amyloid propensity of polyQ is exceptional among the known amyloid-forming sequences on which these predictors were trained and validated, hinting at a very specific nucleation mechanism.

Most pathogenic amyloids feature residues that are hydrophobic and/or have a high propensity for beta strands. Glutamine is neither (Fujiwara et al., 2012; Nacar, 2020; Simm et al., 2016). We reasoned that nucleation therefore corresponds to the formation of a specific tertiary structure unique to Q residues. Structural investigations of predominantly Q-containing amyloid cores reveal an exquisitely ordered tertiary motif wherein columns of Q side chains from each sheet fully extend and interdigitate with those of the opposing sheet, to form a so-called steric zipper (Hervas et al., 2020; Hoop et al., 2016; Schneider et al., 2011; Sikorski and Atkins, 2005). The resulting structural element exhibits extraordinary shape complementarity and an outsized role for very weak (van der Waals) interactions that cumulatively stabilize the structure. Additionally, the terminal amide of each Q side chain in the all-Q steric zipper forms a hydrogen bond to the opposing main chain amide. To minimize confusion while referring to the distinguishing features of all-Q steric zippers, rather than similarly packed but less ordered steric zippers of non-Q residues that lack regular side chain-to-main chain H bonds (Eisenberg and Sawaya, 2017; Sawaya et al., 2007), we will henceforth refer to them as “Q zippers” (**Fig. 2A**).

The high entropic cost of forming an exquisitely ordered Q zipper from the disordered ensemble of polyQ (Chen et al., 2001; Moradi et al., 2012; Newcombe et al., 2018; Vitalis et al., 2007; Wang et al., 2006) stands to be the major contributor to the nucleation barrier. If so, the spacing dependence we identified could be explained by the fact that a single Q zipper would involve only one of the two faces of each beta sheet composed of unbroken beta strands. Because the orientation of successive side chains alternates by 180° along a beta strand, a single Q zipper can be encoded entirely from every other residue in sequence, with the intervening residues having no direct role. Therefore, for short-repeat polypeptides, a Q zipper will be possible only when non-Q side chains occur exclusively after odd-lengthed Q tracts. In contrast, substitutions after even-lengthed tracts would place non-Q side chains on both sides of all potential beta strands and thereby preclude zipper formation (**Fig. 2B**). For both Q_2_X and Q_4_X repeats, steric zippers cannot be formed without including non-glutamine residues. In contrast, even the most substituted odd repeat (Q_1_X) can form a continuous zipper of Qs (**Fig. 2C**).

To determine if the dependence on even/odd spacing and the resulting insertion/exclusion of substituted residues into a Q zipper indeed results from an inability of the zipper to accommodate non-Q side chains, we carried out fully atomistic molecular dynamics simulations with explicit solvent and state-of-the-art force fields to probe the structural stability and conformational dynamics of model polyQ steric zippers. PolyQ amyloids feature antiparallel rather than parallel beta sheets (Matlahov and van der Wel, 2019), and this preference consistently manifests in molecular simulations (Man et al., 2015; Punihaole et al., 2016; Zhang et al., 2016). The simulated amyloid core of polyQ (but not polyN) recapitulates the fully extended, side chain-to-main chain H-bonded steric zipper (Man et al., 2015; Zhang et al., 2016) deduced by structural measurements. We therefore started most simulations as a zipped pair of two-stranded antiparallel β-sheets of 7:Q peptides (model 2O10 (Zhang et al., 2016)). We substituted Q residues in different positions with either asparagine or serine residues, and allowed the structures to evolve for up to 400 ns.

The Q zipper proved to be highly intolerant of non-Q side chains. Substituting any proximal pair of inward pointing Qs led to its dissolution, whereas the structure remained stable when any number of outward-pointing Q residues were substituted **(Fig. 2D, S3A)**. We also ran simulations from parallel beta sheet models that we previously found to be marginally stable for larger sheets (Man et al., 2015; Zhang et al., 2016), but were unable to find any configuration of parallel strands that tolerated substitutions either inside or outside the Q zipper **(Fig. S3B)**, indicating that the nucleating Q zipper cannot be composed of parallel beta-sheets.

Notably, all of the simulated S-containing structures remained intact for longer than the corresponding N-containing structures (not shown), in agreement with the DAmFRET results showing less disruption of nucleation by the former. To gain additional insight into the nucleating structure, we therefore explored potential explanations for the instability imparted by N side chains. As for Q, but not for S, N side chains can simultaneously donate and accept H-bonds. A recent sub-ångström-resolution amyloid microcrystal structure revealed that N side chains can form a pair of H-bonds (termed a “polar clasp”) with adjacent Q side chains on the same side of the strand (i.e. *i* + 2) (Gallagher-Jones et al., 2018). Because this structure is incompatible with interdigitation by opposing Q side chains (**Fig. S3C**), we reasoned that its formation could prevent zipper formation. In order to test this, we quantified the frequencies and durations of side chain-side chain H-bonds in simulated singly N-substituted polyQ strands in explicit solvent, both in the presence and absence of restraints locking the backbone into a beta strand. In short, we found no evidence that Q side chains preferentially H-bond adjacent N side chains (**Fig. S3D,E**), ruling out the possibility of disruptive polar clasps.

The simulations nevertheless did reveal the nature of the destabilizing effect of N side chains. In a perfect Q zipper, the Q side chains form hydrogen bonds both with adjacent side chains and with the backbone of the opposing sheet (**Fig. 2A,E**). When an N side chain falls inside the Q zipper, because it is one methyl group shorter, it cannot make this side chain-to-main chain H-bond. Instead, the side chain of N disrupts the packing of the opposing Q side chains in two steps. First, its terminal amide competes with the adjacent backbone amides for H-bonding with the terminal amide of an opposing Q side-chain, blocking that side chain from fully extending as required for Q zipper stability (**Fig. 2F**). Second, the change in conformation for the detached Q side chain results in steric collisions with adjacent Q side chains. This destabilization propagates through the network of side chains like falling dominos, ultimately unzipping the zipper (**Fig. S3F**).

The side chain of S is shorter than that of N and its terminal group less bulky. Hence it does not intercept H-bonds from the side chains of opposing Q residues (**Fig. S3G**). As a result, the ordering of Q side chains is maintained, and the structure can form, so long as multiple S side chains do not occur in close proximity inside the zipper. Additionally, the shorter side chain of S lessens the entropic barrier to ordered assembly. We suspect this may account for the comparable nucleation frequency of Q_*a*_S constructs relative to pure polyQ **(Fig. 1B, S2C-D)**. In sum, our simulations confirm that Q_(2,4)_N repeats fail to nucleate de novo because the highly ordered structure of Q zippers cannot internally accommodate non-Q side chains in general, and N side chains in particular. Therefore, the conformational nucleus for “polyQ” amyloids is a Q zipper that can only be formed by segments of sequence with unilaterally contiguous Qs.

### Bilaterally contiguous glutamines encode shorter, laminated zippers

We observed that the dependence on spacing for amyloid formation declined sharply beyond 5 consecutive Qs. Above this apparent threshold, nucleation occurred de novo irrespective of the spacing of non-Q residues **(Fig. 3A, S2B)**, suggesting the onset of an alternative nucleating structure composed of strands extending approximately six residues. The fraction of cells transitioning to the high-FRET state increased both with the expression level and consecutive Q length above this threshold (**Fig. 3A, Fig. S2B**).

**Figure 3.**
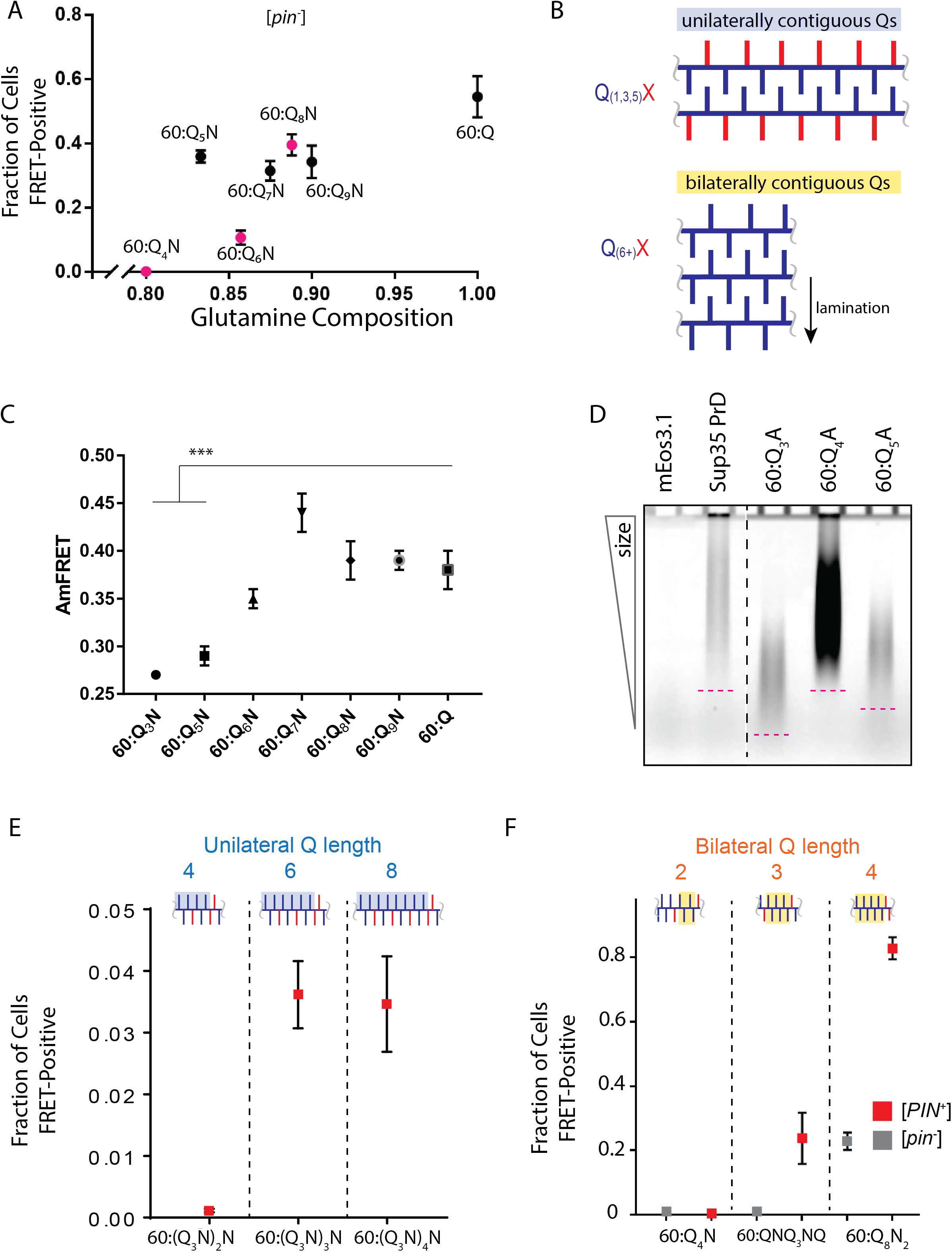
Bilaterally contiguous glutamines encode shorter, laminated zippers. A. Scatter plot of the fraction of [*pin*^-^] cells with amyloid versus the fraction of glutamines in 60:Q_a_N proteins, showing that for even values of *a* (red dots), nucleation only occurs for *a* ≥ 6. Shown are means +/- SD of quadruplicates. B. Schematic of single Q zipper and laminated Q zipper architectures. C. AmFRET values of [*pin*^-^] cells containing amyloids of the indicated sequence variants. Shown are means +/- SD of the median AmFRET values of quadruplicates. *** p < 0.001, t-test. D. SDD-AGE characterizing amyloid length distributions for mEos3.1-tagged 60:Q_3_A, Q_4_A and Q_5_A polypeptides. Note that the lower boundary of the amyloid smear is lower for Q_3_A and Q_5_A relative to Q_4_A and Sup35 PrD, a typical Q/N-rich amyloid. Data are representative of multiple experiments. E. Fraction of [*PIN^+^*] cells with amyloid for 60:Q_3_N-derived sequences. Sequences are frame-shifted to give unilaterally contiguous lengths of 4, 6 or 8 Qs. Inset shows side-chain arrangements as in 3A. Shown are means +/- SD of quadruplicates. F. Fraction of [*PIN^+^*] or [*pin*^-^] cells with amyloid for 60:Q_4_N-derived sequences. Sequences are frame-shifted to give bilaterally contiguous lengths of 2, 3, or 4 Qs (i.e. on both sides of the beta strand). Inset shows side-chain arrangements as in 3A. Shown are means +/- SD of quadruplicates.

Because steric zipper ordering on one side of a beta strand configures the backbone for steric zipper ordering on the other side of the strand, provided both sides have contiguous Qs, the acquisition of an alternative amyloid structure with increased density by Q_(4,6+)_ repeats could be explained by the formation of short Q zippers on *both* sides of the beta strands. This would result in a laminated architecture, with multiple layers of beta sheets bonded by Q zippers (**Fig. 3B**).

Consistent with this interpretation, we observed that cells containing Q_(6+)_ amyloids reached higher AmFRET than those containing Q_(3,5)_ amyloids (**Fig. 3C**, **Fig. S2B-D**), suggesting that the former amyloids have a higher density (number of subunits in cross-section) than those of the latter.

As a more direct probe of structural differences between the two classes of Q zipper amyloids, we exploited the resistance of amyloids to denaturation in strong detergents (Nielsen et al., 2007; Villali et al., 2020), to compare their distributions of lengths using semi-denaturing detergent-agarose gel electrophoresis (SDD-AGE) (Halfmann and Lindquist, 2008; Kryndushkin et al., 2003). We observed a dramatic and consistent dependence of amyloid lengths on Q contiguity (**Fig. 3D, S4A**). Specifically, Q_3_ amyloids ran the lowest, followed by Q_5_. Q_(4,6+)_ repeats ran the highest. These differences cannot be explained by different elongation rates of the corresponding amyloids, because amyloid formation by the former achieves steady state more rapidly than the latter. Nor are they consistent with different primary nucleation rates (Knowles et al., 2009), given the extended region of bimodal DAmFRET for the former, which implies a greater barrier to primary nucleation. Therefore, the length differences must be attributed to a secondary process(es) such as fragmentation. Based on established relationships of fragmentation to structure (Tanaka et al., 2006; Verges et al., 2011), the much shorter lengths of Q_(3,5)_ amyloids suggest a thinner, more fragile architecture than those of Q_(4,6+)_, consistent with lamination of the latter.

To investigate the lengths of continuously ordered beta strands in each type of zipper, we designed variants of 60:Q_3_N and 60:Q_4_N with differing lengths of unilateral or contralateral contiguity, respectively. We found that in the absence of contralateral Qs, de novo amyloid formation retained a low concentration-dependence only for six or more unilaterally contiguous Qs **(Fig. 3E, S4B)**. In contrast, only three unilaterally contiguous Qs were required in the presence of contralateral Qs **(Fig. 3F, S2B,S4C)**. These findings imply that the beta strands in the single Q zipper nucleus extend for 11-12 residues, while those in the laminated Q zipper nucleus extend for only 6 residues. This latter number is consistent with the ideal lengths of antiparallel strands (Stanger et al., 2001), and is remarkably close to the critical strand length of 7 that had been deduced for apparently monomeric nucleation in vitro (Thakur and Wetzel, 2002). The slightly longer threshold in vitro can be attributed to charge repulsion from the lysine pair that had been appended to both ends of the polyQ tract (Walters and Murphy, 2009). Our constructs lack this element, as do endogenous pathologic polyQ tracts.

The different strand lengths required to form single or laminated Q zipper nuclei explain why 60:Q_4_N cannot form either type of nucleus. This is illustrated by comparing the three polypeptides with identical length, composition, and unilateral contiguity (60; 48 Qs, 12 Ns; and 4, respectively). In the case where contralateral zipper-forming frames were maximally offset (60:Q_4_N), amyloid did not form. Shifting the registry by one position (60:QNQ_6_NQ) allowed for very sporadic amyloid formation only at very high concentrations, and shifting the registry by one more position (60:Q_8_N_2_) -- to maximize overlap between contralateral zippers -- further increased amyloid formation **(Fig. 3F, S2B,S4C)**. We conclude that the extraordinary resilience of Q_4_N against amyloid formation can be attributed to its being the highest possible density of Qs that still lacks sufficient contiguity both unilaterally and bilaterally.

### Q zipper nucleation occurs within monomers

Another intriguing difference between unilaterally and bilaterally contiguous sequences is that the former nucleated with a greater efficiency at low concentrations (**Fig. 4A**). For example, the fraction of [*pin*^-^] cells in which 60:Q_5_N formed amyloid rose sharply with expression up to 10^7^ AFU, where it plateaued at approximately 25%. In contrast, the fraction of cells in which 60:Q_(6+)_N formed amyloid rose steadily with expression even beyond 10^8^ AFU. We were able to envision two, not mutually exclusive mechanisms whereby nucleation could be compromised at high concentrations. First, there may exist within the cell a limiting amount of some nucleation-promoting factor, such as a molecular chaperone. As the concentration of the protein of interest increases, this factor becomes saturated. Second, the protein may partition into “off-pathway” oligomers or condensates at high concentration, within which the nucleating conformation is disfavored.

**Figure 4.**
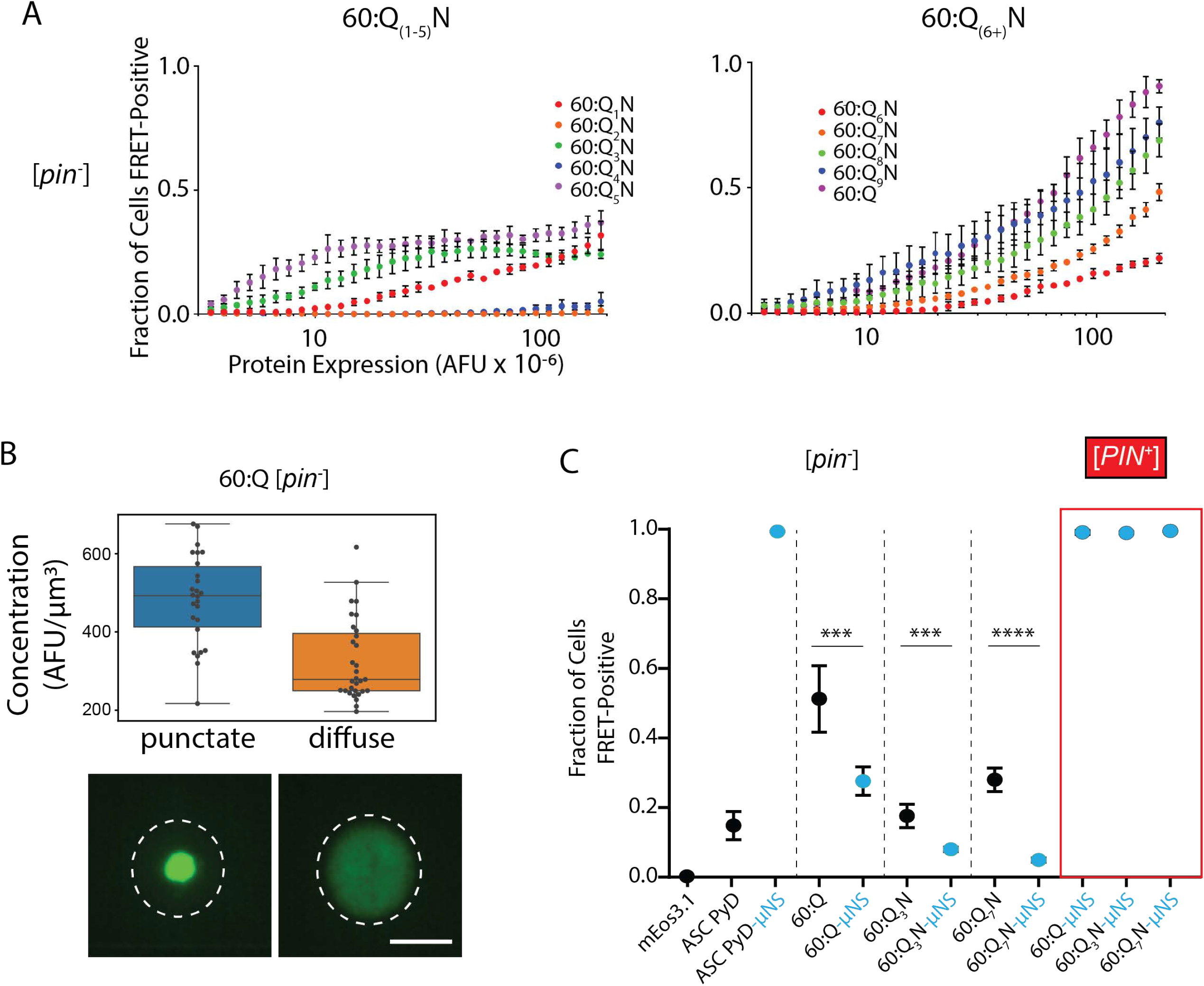
Q zipper nucleation occurs within monomers. A. Fraction of cells AmFRET-positive as a function of expression level, for [*pin*^-^] cells expressing 60:Q_(1-5)_ (left) or 60:Q_(6+)_ (right). Note the plateau in 60:Q_(3,5)_ profiles suggestive of inhibitory oligomerization. B. Distribution of cytosolic concentrations (AFU/μm^3^) of 60:Q-mEos3.1 expressed in [*pin*^-^] cells either lacking or containing puncta, showing that the protein remains diffuse even when supersaturated relative to amyloid. Representative diffuse or punctate cells (N = 31 and 26, respectively) of equivalent total concentration are shown. Scale bar: 5 μm. C. Fraction of cells with amyloid for the indicated proteins expressed with or without genetic fusion to μNS. Artificial condensation sufficed to drive nucleation for all cells expressing ASC PyD, which has an oligomeric nucleus. It likewise did so for 60:Q, 60:Q_3_N, and 60:Q_7_N specifically in [*PIN*^+^] cells, whereas it inhibited their nucleation in [*pin*^-^] cells. Shown are means +/- SD of triplicates. *** p < 0.001, **** p < 0.0001; t-test.

PolyQ has been observed to form non-amyloid oligomers and condensates, which has led to the prevailing paradigm that phase separation precedes and facilitates the nucleation of conformational order (Borcherds et al., 2021; Camino et al., 2021; Crick et al., 2013, 2006; Dignon et al., 2020; Fisher et al., 2021; Halfmann et al., 2011; Peskett et al., 2018; Posey et al., 2018; Vitalis and Pappu, 2011; Yang and Yang, 2020). The second explanation above would seem to contradict this paradigm. To investigate whether phase separation may be occurring in our system, we inspected the subcellular localization of representative proteins in different regions of their respective DAmFRET plots in both [*pin*^-^] and [*PIN*^+^] cells. Using imaging flow cytometry, we found that all high AmFRET cells contained large round or stellate puncta, as expected for amyloid deposition, whereas virtually all cells with low or no AmFRET contained entirely diffuse protein, even at high expression in a bimodal regime (**Fig. S5A**). Next, we used confocal fluorescence microscopy to compare total cytosolic concentrations of 60:Q expressed in [*pin*^-^] cells that either lacked or contained puncta, and found that the distributions of concentrations overlapped (**Fig. 4B**), suggesting the protein remained dispersed prior to amyloid formation even when deeply supersaturated. We conclude that polyQ, even when expressed to high micromolar concentrations, does not visibly phase separate (as detectable by fluorescence microscopy) prior to amyloid formation in living cells.

We next probed the role of oligomerization in amyloid formation, by relaxing the dependence of oligomerization on protein expression level. To do so, we lowered solubility by expressing select proteins as fusions to a well-characterized modular moiety, μNS, that forms coiled coil-mediated multimers at low concentration and does not itself form amyloid (Brandariz-Nuñez et al., 2010; Kroschwald et al., 2015; Schmitz et al., 2009). As a control for an oligomeric nucleus, we used the human inflammasome scaffold protein, ASC. The pyrin domain of ASC self-associates via electrostatic interfaces into a monomorphic three-stranded polymer (Lu et al., 2014) that is physiologically rate limited by nucleation (Cai et al., 2014; Khan et al., 2018). We found that genetically fusing μNS to ASC pyrin drove it to nucleate in every [*pin*^-^] cell, as expected (**Fig. 4C, S5B**). Note that amyloid formation by μNS-fused proteins results in a higher AmFRET value than that of μNS condensates themselves, and the latter does not depend on fusion partner (**Fig. S5B-C**, and other data not shown). Note also that amyloids of fusion proteins tend to have lower AmFRET than that of the corresponding non-fused proteins; we attribute this to the fact that larger proteins occupy more volume per fluorophore unit. We next asked how μNS-driven condensation influenced nucleation for Q zipper amyloids. In striking contrast to ASC pyrin, representative QaX proteins fused to μNS nucleated de novo at *reduced* frequencies to their unfused counterparts (**Fig. 4C, S5B**). This effect was limited to the nucleation step, as the μNS-fusions went on to populate a uniform high-FRET state in every cell that contained [*PIN*^+^]. That is, fusion-mediated oligomerization appears to have driven to amyloid the kinetically arrested species that had previously populated the low arm of the sigmoid. To confirm that the reduction in nucleation resulted from the protein’s local condensation rather than a specific effect of μNS, we also performed experiments with a separate fusion partner, human ferritin heavy chain, which forms discrete 24-mer homo-oligomers (Bracha et al., 2018). Again, nucleation was reduced in [*pin*^-^], specifically, relative to non-fused proteins (not shown). These results reveal that oligomerization of disordered polyQ impairs the rate-limiting conformational fluctuation required for it to acquire a minimal Q zipper. Once that zipper has nucleated, however, oligomerization drives it toward amyloid. In other words, the sequence-encoded homogeneous nucleus contains an exclusively intramolecular Q zipper.

### Lamination competes with Q zipper lengthening

Q_(6+)_ repeats also recapitulated the susceptibility of pure polyQ to a conformational template. Whereas [*PIN*^+^] cells expressing Q_(3,5)_ polypeptides populated both high- and low-AmFRET values over a large window of expression, those expressing Q_(6+)_ populated a much narrower range of AmFRET values for a given expression level **(Fig. S2B-D)**. These data suggest that lamination specifically reduces a critical density fluctuation, which is more limiting in the presence of [*PIN*^+^]. In other words, lamination stabilizes oligomers with relatively unstable Q zippers, and this compensates for imperfections in the Q zipper that must occur at the interface of polyQ and the [*PIN*^+^] amyloid template. It also explains why lamination lowers the strand length requirement. In support of this interpretation, we observed that Q_(6+)_ transitioned from low- to high-AmFRET in a continuous, sigmoidal fashion, indicating that elongation or maturation of the nascent amyloids, rather than nucleation per se, had become limiting. Continuous DAmFRET transitions can be interpreted as the formation of a distribution of multimers wherein the extent of assembly, in terms of both amount and order, increases with expression level (Posey et al., 2021). To identify trends between AmFRET distributions that lack bimodal regimes, we plotted the mean of AmFRET values in serial bins of protein expression level (**Fig. 5A**). The lower arms of the resulting curves represent cells with low but non-negligible levels of AmFRET. We observed that, as the consecutive Q length increased, the midpoint of the distribution shifted to lower expression levels (**Fig. 5A**).

**Figure 5.**
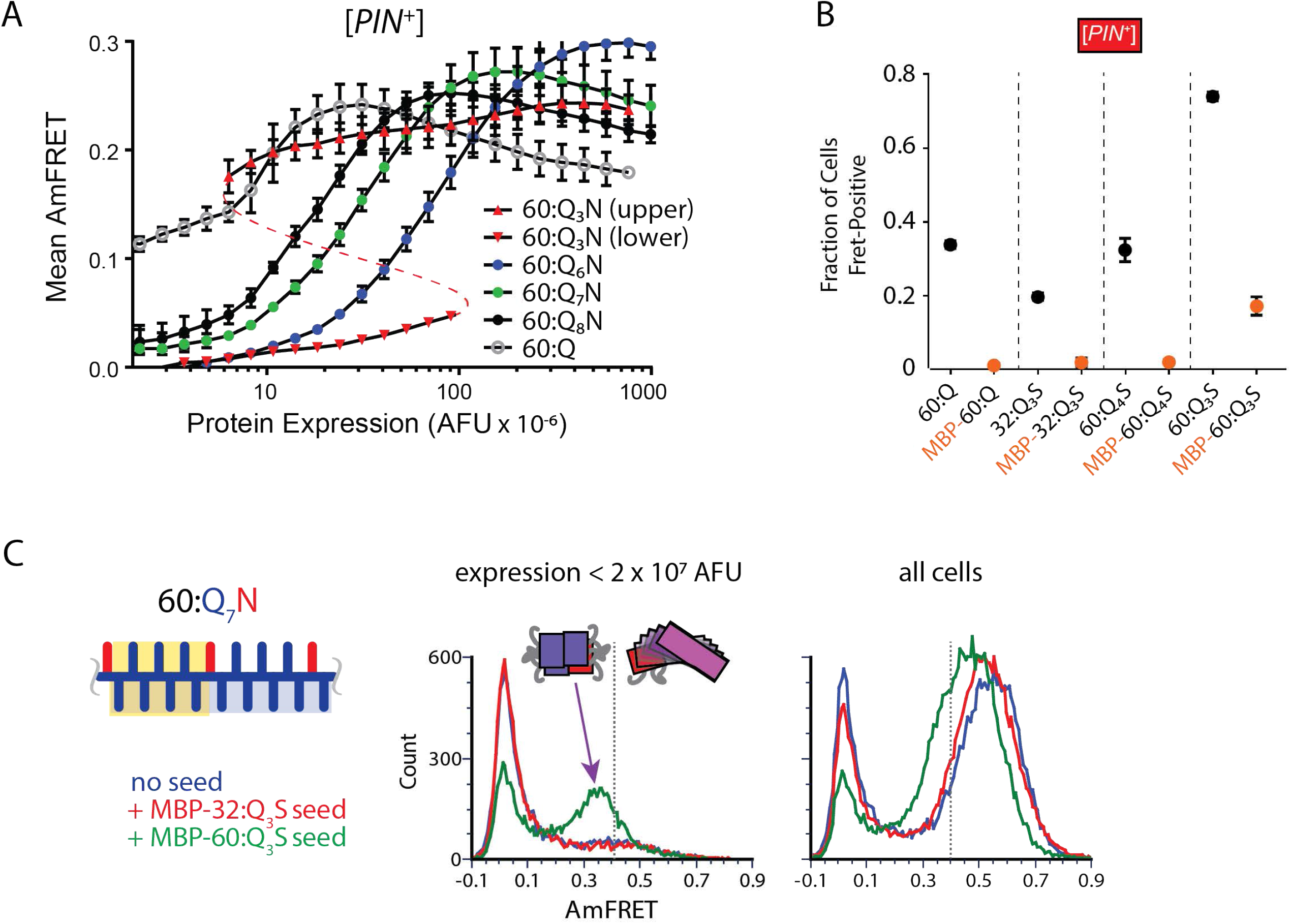
Q zipper nucleation occurs within monomers. A. Graph of spline fits of AmFRET values for 60:Q_(6+)_N and the upper and lower populations of 60:Q_3_N, in [*PIN^+^*] cells, showing that Q_(6+)_N but not Q_3_N have appreciable AmFRET at low expression, that the Q_(6+)_N AmFRET level is lower than that of the upper Q_3_N population at low expression, and higher than that of the upper Q_3_N population at high expression. The red dashed line denotes that the upper and lower traces of 60:Q_3_N represent subpopulations of the same samples. Shown are means +/- SD of quadruplicates. B. Fraction of [*pin*^-^] cells with amyloid for the indicated variants either lacking or containing a genetic fusion to MBP. C. Histograms of AmFRET values of 60:Q_7_N at low expression (<2×10^7^ AFU) showing the appearance of a mid-FRET population, indicative of single Q zipper amyloids, only when seeded by MBP-60:Q_3_S.

Despite the reduced nucleation barrier for Q_(6+)_, the level of AmFRET achieved at low expression (e.g. < 10^8^ AFU for 60:Q_6_N and < 10^7^ AFU for Q:60), was reduced relative to Q_(3,5)_ amyloids (**Fig. 5A, S2B-D**), suggesting that the laminated oligomers are both off-pathway to, and preclude the formation of, long Q zipper species. That their formation is nucleation-limited means that this inhibitory activity cannot be attributed simply to a depletion of naive monomers. Rather, they must share a critical intermediate with long Q zipper amyloid. This intermediate is presumably the structural element common to both of them, namely, a single short Q zipper. This short zipper, once formed, can *either* extend to form amyloids, or laminate to stabilize “pre-amyloid” oligomers.

To explore this notion, we destabilized oligomers by expressing proteins as fusions to a well-established solubilizing moiety -- maltose binding protein (MBP). MBP does not interfere with the folding of its fusion partner but, due to its bulk, sterically limits its access to other protein molecules (Raran-Kurussi and Waugh, 2012). Fusion to MBP should therefore amplify the density component of nucleation barriers, rendering nucleation more concentration-dependent. It will also restrict the dimensions of self-assemblies, resulting in lower AmFRET. Finally and most importantly, it should bias amyloid conformers toward longer Q zippers.

To validate the approach, we compared DAmFRET statistics for nucleation by ASC, without and with fusion to MBP. The C_min_ statistic theoretically corresponds to the phase boundary; the EC50 statistic represents the concentration at which nucleation has occurred in half the cells, and the □ (delta) statistic describes the independence of nucleation on concentration, where conformationally-limited nucleation has higher □ values (Khan et al., 2018). As expected, we found that MBP increased both C_min_ and EC_50_, but did not change □ (**Fig. S6A**). It also reduced AmFRET levels by approximately 30%. We then tested the effect of MBP on 60:Q as well as Q_3_S polypeptides below or above the length threshold (32 and 60, as determined below) for intramolecular Q zipper nucleation. We found that MBP completely eliminated detectable assemblies for 60:Q and 32:Q_3_S. In contrast, MBP-60:Q_3_S retained a bimodal distribution, albeit with increased concentration-dependence and reduced AmFRET of the nucleated assemblies (**Fig. 5B, S6B**). To minimize any confounding effect of MBP on oligomerization post nucleation that may preclude a detectable increase in AmFRET, we also performed DAmFRET on an unmodified 60:Q_3_N expressed in trans, to report on nucleation occurring in non-fluorescent versions of the above MBP fusions. As expected, MBP-60:Q_3_S, but not MBP-32:Q_3_S nor MBP-60:Q, promoted 60:Q_3_N amyloid formation **(Fig. S6C)**. Together, these results confirm that lamination competes with Q zipper lengthening en route to mature polyQ amyloid, and that oligomerization relaxes that competition.

Finally, we probed the effect of MBP on the switch between single and laminated Q zipper preferences that occurs at Q_7_N. This sequence represents the minimal consecutive Q length with continuous unilateral contiguity that nevertheless exhibited the continuous sigmoidal DAmFRET of laminated Q zipper-forming sequences. That 60:Q_7_N fails to nucleate single zipper amyloids suggests that the alternative, laminated Q zipper has a lower nucleation barrier and therefore involves a less extreme conformational fluctuation. If so, the ability of Q_7_N to nucleate a single zipper amyloid should be exposed either by selectively rewarding conformational fluctuations -- by expressing it in the presence of orthogonal long zipper amyloids. We therefore expressed 60:Q_7_N in [*PIN*^+^] cells that also expressed nonfluorescent MBP fusions of 32:Q_3_S or 60:Q_3_S in trans. MBP-32:Q_3_S had no effect on 60:Q_7_N. In contrast, MBP-60:Q_3_S allowed the cells to populate *both* the intermediate- and high-AmFRET states (**Fig. 5C, S6C**), indicative of single and laminated Q zipper amyloids, respectively. Importantly, low expression of 60:Q_7_N favored the former, while high expression favored the latter, consistent with their relative dependencies on oligomerization. We conclude that the MBP-60:Q_3_S seeds unmasked a latent ability of 60:Q_7_N to self-assemble (once nucleated) into amyloid with a single Q zipper structure. This result confirms that 60:Q_7_N can form either type of amyloid, but is kinetically prevented from nucleating long Q zippers de novo due to the relative ease with which short Q zippers are stabilized through lamination.

Altogether, our experiments to manipulate critical fluctuations in density and conformation reveal the existence of a bifurcated pathway from polyQ monomer to amyloid. Specifically, the critical conformational fluctuation occurs within a disordered monomer to produce a minimal intramolecular Q zipper, which can then either laminate or lengthen in a manner governed by the presence or absence, respectively, of contralateral Qs (**Fig. 6A**). Low concentrations and solubilization favor assemblies with longer strands and fewer sheets, while high concentrations and condensation favor assemblies with shorter strands and more sheets (**Fig. 6B**). The former manifest with reduced AmFRET levels and polymer lengths, relative to the latter. [*PIN*^+^] accelerates the rate-limiting monomer nucleation step and thereby amplifies these differences at the population level. The proposed pathway explains multiple features of our DAmFRET and SDD-AGE data not yet discussed: while Q_(3,5)_X polypeptides nucleated the predominant lower-AmFRET form at low concentrations, a higher AmFRET form consistent with laminated amyloid emerged at high concentrations (for example, compare expression levels greater than, versus less than, 10^8^ AFU for 60:Q_5_N and 60:Q_3_S in **Fig. S2B** and **S2C**, respectively). [*PIN*^+^] inhibited this transition (**Fig. S2B-D**), and reduced the lengths of SDS-resistant species (**Fig. S4A**).

**Figure 6.**
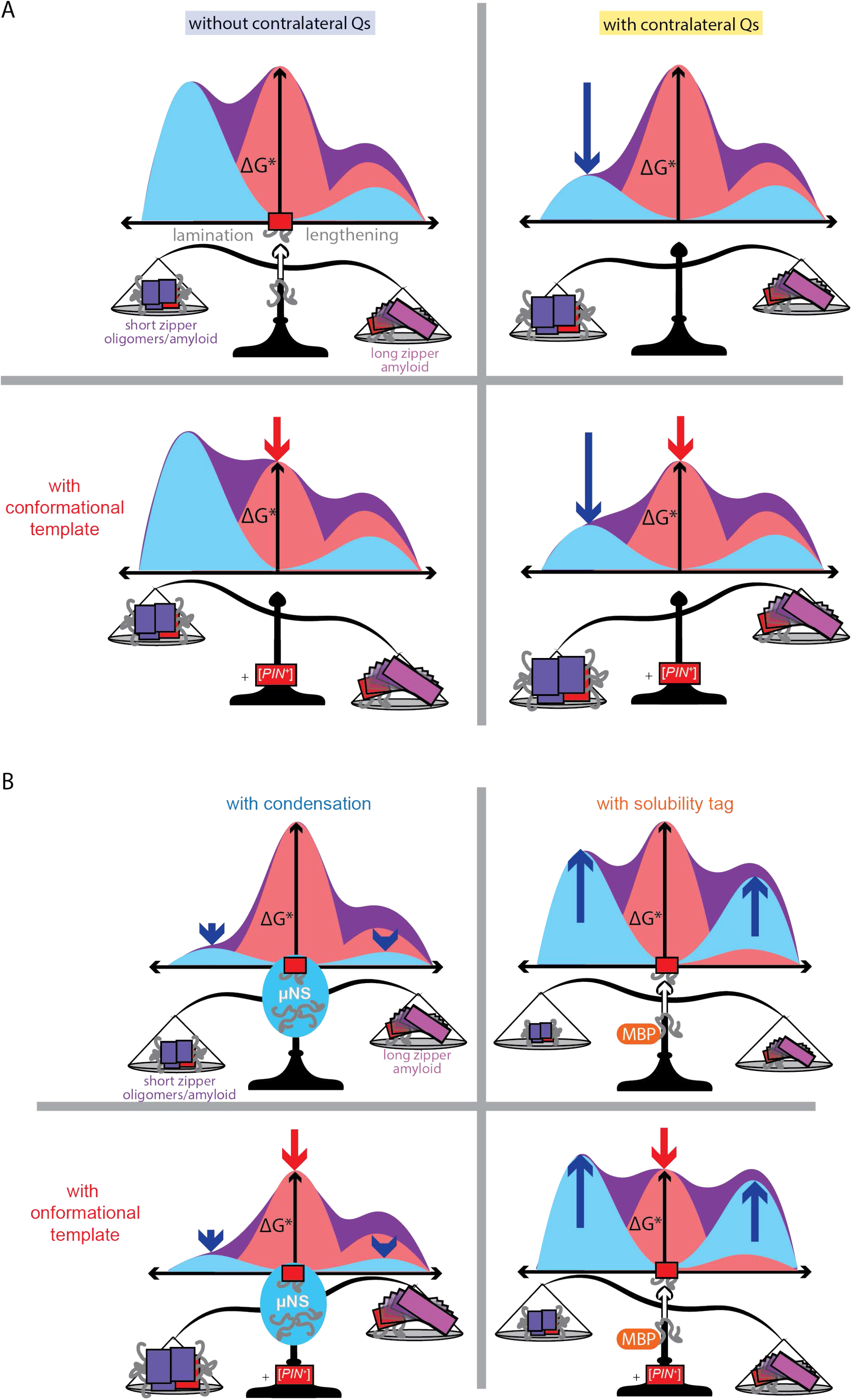
Reaction coordinate diagrams for a bifurcated amyloid nucleation pathway. A. Reaction coordinate diagrams for heterogeneous amyloid nucleation, whereby the disordered monomer acquires a minimal Q zipper (red rectangle), whose subsequent oligomerization extends the Q zipper either by lamination or lengthening, to produce short zipper amyloid-like oligomers or long zipper amyloids. The former transition is promoted by density fluctuations and contralateral Qs. Providing a conformational template, in the form of [*PIN^+^*] (bottom row), accelerates Q zipper nucleation and consequently amplifies these preferences. B. Decreasing solubility by genetic fusion to μNS, does not accelerate Q zipper nucleation (top left), but does accelerate the maturation of a pre-existing Q zipper to amyloid, analogous to nucleation in [*PIN^+^*] cells (bottom left). Increasing solubility by genetic fusion to MBP decelerates Q zipper lamination (top right), and this effect is most pronounced in [*PIN*^+^] cells (bottom right).

### The Q zipper nucleus exhibits the pathologic length threshold

If the rate-limiting Q zipper indeed occurs within a single polypeptide, nucleation will become sharply more concentration-dependent for sequences that are too short to form that element (**Fig. 7A**). To evaluate potential length thresholds for single Q zipper nuclei, we generated a series of length variants for Q_3_S (S was chosen for its relatively modest nucleation defect that afforded a wider window of informative length variants) and, for each, we evaluated the dependence of nucleation on concentration by comparing the EC_50_ and □ statistics derived from DAmFRET (**Table S3**). We observed a positive linear relationship between □ and EC_50_, as expected, as the length of the polypeptide fell from 40 to 36 (**Fig. 7B, S7A**). The relationship became negative below 36, implying an increasing dependence of nucleation on intermolecular interactions. We interpret this value as an approximation of the minimum length of polypeptide, in the absence of other interacting domains, required for a Q zipper to form within a single polypeptide.

**Figure 7.**
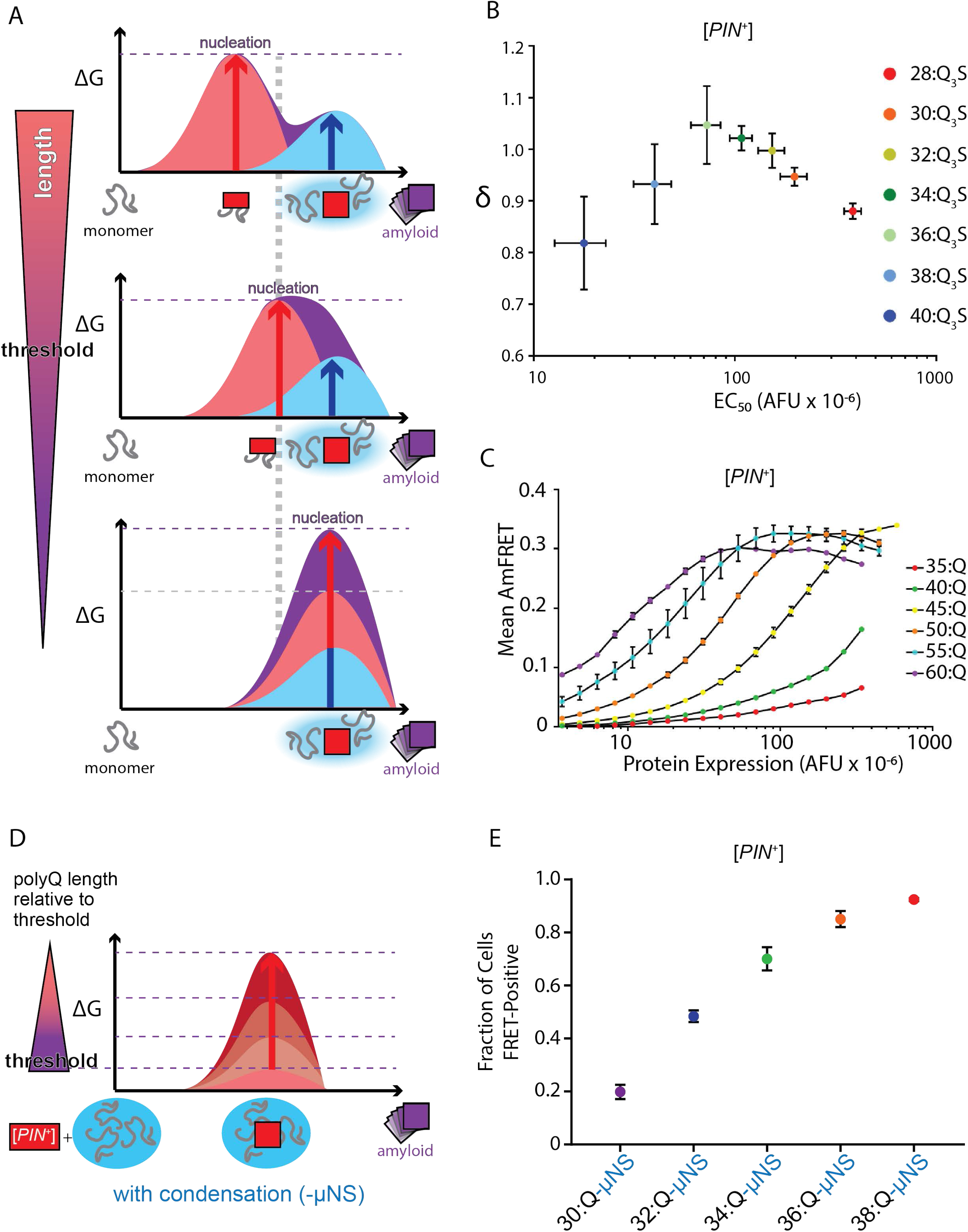
The Q zipper nucleus exhibits the pathologic length threshold. A. Reaction coordinate diagram schematizing the relationship of polypeptide length to heterogeneous amyloid nucleation barriers. As length decreases below that required to nucleate intramolecularly, the nucleation barrier will increase because the limiting conformational fluctuation now becomes dependent on a density fluctuation. B. Scatter plot of the concentration-dependence of nucleation (delta) versus the concentration at which the protein has nucleated in half of the cells (EC50), obtained from Weibull fits of DAmFRET profiles of different length variants of Q_3_S expressed in [*PIN^+^*] cells. Shown are means +/- SD of quadruplicates. C. Spline fits of DAmFRET data for polyQ length variants 35 and longer expressed in [*PIN^+^*] cells. D. Reaction coordinate diagram schematizing the relationship of polyQ length to the nucleating conformational fluctuation. The proteins are fused to μNS to eliminate density fluctuations from the nucleation barrier, and expressed in [*PIN^+^*] cells to achieve detectable nucleation frequencies. The conformational energy barrier increases, and acquires a darker shade of red, with shorter lengths of polyQ. E. Fraction of [*PIN^+^*] cells with amyloid for the indicated polyQ length variants fused to μNS.

We next explored the length-dependence for amyloid formation by uninterrupted polyQ. We found that amyloid failed to form below 35:Q, formed to an almost imperceptible extent at 35:Q, and formed robustly at 40:Q and above **(Fig. 7C, S7B-D)**. The sigmoidal transition from low to high AmFRET in [*PIN*^+^] cells shifted to lower concentrations with increasing polypeptide length. Given the inclination of polyQ toward metastable intermediates, this relationship suggests an accelerated progression of laminated zippers to amyloid by longer polyQ. Note, however, that the continuous nature of the transition from low to high AmFRET in this case means that these results cannot be interpreted with respect to a threshold length for the nucleus itself. In other words, we cannot exclude from this experiment that nucleation may occur below 35:Q and simply fail to elongate sufficiently over the accessible concentration range to produce detectable AmFRET.

Therefore, to directly interrogate the length threshold for the polyQ amyloid nucleus itself, we needed to eliminate the concentration-dependence of laminated Q zipper maturation. To do so, we pre-condensed the polyQ length variants to high local concentration by expressing them as fusions to μNS in [*PIN*^+^] cells (**Fig. 7D**). By partitioning the proteins into a condensate, this fusion minimizes any contribution of polyQ length to density heterogeneities. We found that polyQ nucleated in almost all cells at lengths 38 and above, but in a decreasing fraction of cells at lengths 36 and below (**Fig. 7E, S7E**). These data suggest that Q zipper formation is compromised when the continuous chain length falls below 36, which provides a lower bound on the length threshold of an intramolecular Q zipper.

### Structural model of the polyQ amyloid nucleus

Collectively, our data constitute an empirical survey of relevant points on the energy landscape of polyQ amyloid formation, which we illustrate in **Fig. 8**. Excursions from naive disordered monomers, represented by the basin at bottom left, are described in terms of intermolecular density and intramolecular order. Mature amyloids featuring lengthened and laminated Q zippers lie in a deep well in the opposite corner from the disordered monomers. A rare conformational fluctuation within a monomer (dashed red arrow) creates an embryonic Q zipper.

**Figure 8.**
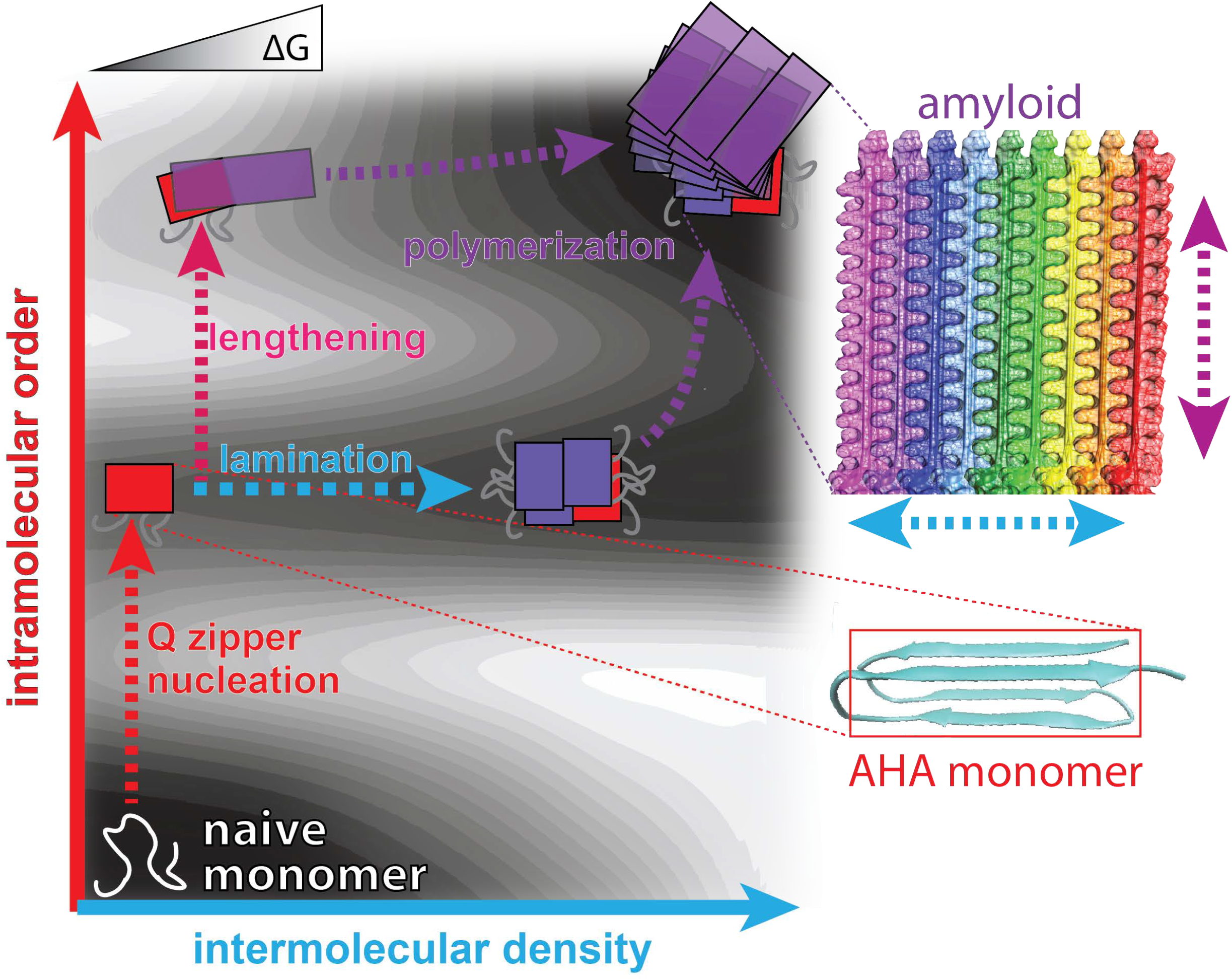
Structural model of the polyQ amyloid nucleus. The hypothetical surface of free energy in the plane of intramolecular (Q zipper length, red axis) and intermolecular order (density, blue axis) in a supersaturated solution of polyQ molecules. Qualitative topological features of the landscape, but not absolute heights and positions, are as deduced herein. Naive monomers exist in a local energy minimum at maximum disorder, while mature amyloid exists in a global energy minimum with long laminated Q zippers (image from Boatz et al. 2020, with permission). The route from monomer to amyloid proceeds through either of two horizontal basins. The lower basin represents short Q zippers and features the nucleating AHA monomer at the shallow end. The basin descends gradually toward amyloid via lamination-driven oligomerization. The upper basin represents long Q zippers, and descends more steeply toward amyloid due to the increased intermolecular connectivity of long zippers. Transitions from the lower to upper basin involve high-energy lengthening of short Q zippers.

Do the constraints identified here offer a glimpse of the embryonic Q zipper structure? Our data indicate that the critical Q zipper can be accessed by polypeptides as short as approximately 36 residues and has strands of length 6. Allowing for connecting turns and arcs of two to four residues each (Chou and Fasman, 1977; Hennetin et al., 2006), it can contain no more than four β-strands. If we further grant from pioneering in vitro experiments (Bhattacharyya et al., 2005; Buchanan et al., 2014; Chen et al., 2002; Kar et al., 2013, 2011; Sikorski and Atkins, 2005) that the formation of a hairpin -- and not an arch -- is rate-limiting, then the four strands are connected by a single turn and two arcs. This allows for only two possible non-equivalent Q zipper configurations. One has the first and fourth strands unpaired and flanking either side of the central hairpin. We simulated this configuration and found it to be extremely unstable (**Fig. S8A**) and therefore discarded it from further consideration. The other configuration has the first and fourth strands paired and flanking one side of the central hairpin (**Fig. S8B**). This hypothetical fold of polyQ, along with the alternative configuration having two hairpins and one arch, had previously been simulated by multiple groups and found to be more stable than any single sheet or parallel stranded zipper configuration (Man et al., 2015; Miettinen et al., 2012). We therefore conclude that the nucleating monomer most likely comprises a pair of two-stranded antiparallel sheets in a super-secondary topology of β-(arch, hairpin, arch), and will be hereafter referred to as “AHA”.

Our observations suggest that the AHA monomer does not appreciably polymerize, and so is not yet “amyloid”. We suspect the axial surface is too short or unstable to serve as an intermolecular template for converting the conformations of new subunits, a process that for polyQ takes considerably longer than the rate of naive molecules encountering the surface (Bhattacharyya et al., 2005). Our results imply that the emergence of an amyloid axis of polymerization requires that the Q zipper first grow in either of two orthogonal directions to produce a larger and more stable templating surface. This occurs via oligomerization coupled to either lamination (**Fig. 8**, dashed blue arrow), or lengthening (**Fig. 8**, dashed fuschia arrow). The latter is associated with a larger nucleation barrier and is therefore a more extreme conformational fluctuation. Consequently, early Q zipper growth proceeds through lamination, resulting in an accumulation of amyloid-like oligomers. Subsequent iterations of subunit addition present new opportunities for the Q zipper to lengthen and reinforce the axis of polymerization. Consequently, our model predicts a directional progression during polymerization (dashed purple arrows) toward multiply laminated fibers with fully extended hairpins, consistent with the latest models for mature polyQ amyloids (Boatz et al., 2020).

To understand the nature of the second energy barrier associated with Q zipper lengthening, we used the constraints derived herein to create a structural model for the minimal hypothetical subunit of long zipper amyloid. Our data imply that this species can be accessed by polypeptides as short as approximately 36 residues and has unilaterally contiguous strands of length 11. It therefore contains three beta strands connected by a single turn and arc. We performed molecular simulations to investigate the stability of this structure as an isolated monomer and found that it fell apart within 200 ns (**Fig. S8C**). Together with the above simulation, this result reveals that a Q zipper cannot be maintained by just three strands even when those strands are the length required for long Q zipper amyloid. We then asked if it persisted for longer when joined to a second such conformer across the hypothetical axial interface. Indeed, the resulting homodimer not only persisted, but became slightly *more* ordered over the duration of the simulation (**Fig S8D**), consistent with the onset of energetically downhill polymerization.

## Discussion

### PolyQ amyloid begins *within* a molecule

The initiating molecular event leading to pathogenic aggregates is the most important yet unknown and difficult to study step in the progression of age-associated neurodegenerative diseases including polyglutaminopathies such as Huntington’s Disease. Here we used our recently developed assay to identify sequence features that govern the dependence of nucleation frequency on concentration and conformational templates, to deduce that amyloid formation by pathologically expanded polyQ begins with the formation of a minimal Q zipper within a single polypeptide molecule.

Kinetic studies performed both with synthetic peptides in vitro (Bhattacharyya et al., 2005; Chen et al., 2002; Kar et al., 2011) and recombinant protein in animals (Sinnige et al., 2021) and neuronal cell culture (Colby et al., 2006) have found that polyQ aggregates with a reaction order of approximately one. This has been interpreted as evidence that pathogenic nucleation occurs within a monomer (Bhattacharyya et al., 2005; Chen et al., 2002; Kar et al., 2011). However, this interpretation assumes homogeneous nucleation, which is invalidated by observations of non-amyloid multimers prior to amyloids in vitro (Crick et al., 2006; Vitalis and Pappu, 2011). We found that even when accounting for heterogeneous nucleation, the original conclusion of Chen et al. holds. Specifically, the rate-limiting step for amyloid formation from soluble monomers *does* occur within a monomer, and the relevant multimeric heterogeneities *follow* rather than precede that event. Because they interact via the defining secondary and tertiary elements of “amyloid” conformation, these “pre-amyloid” oligomers are not condensates in the sense of first order phase separation. Rather, they are equivalent to extremely early amyloid polymorphisms that have not yet achieved a dominant axis of polymerization.

Given that monomers can nucleate, and that there are thousands of nucleation-competent monomers in each neuron of a patient’s brain, why does aggregation and disease onset still take years? The reason lies in the peculiar properties of polyglutamine. Glutamine residues lack a strong preference for any particular secondary structure and strongly solvate the polypeptide backbone. Hence, polyglutamine populates a collapsed ensemble of disordered states (Crick et al., 2006; Kang et al., 2017; Walters and Murphy, 2009). The Q zipper is plausibly the only stable structure that can be formed by polyQ, as it satisfies the side chain H-bonding potential without disrupting backbone H-bonding. Critically, the strong tendency toward disorder for polyQ persists even when the entropic penalties for beta structure have been prepaid (Vitalis and Pappu, 2011), suggesting that it is the formation of the defining tertiary element of polyQ amyloid, i.e. the Q zipper, rather than the beta sheets themselves, that imposes the entropic bottleneck narrow enough to delay nucleation over the timescales of human lifespan.

Pioneering studies revealed that the nucleation event involves the formation of a beta hairpin (Bhattacharyya et al., 2005; Chen et al., 2002; Kar et al., 2013, 2011; Sikorski and Atkins, 2005). In our model, and consistent with early proposals by Wetzel and colleagues (Kar et al., 2011), the termini of the hairpin fold back toward the beta turn. Unlike other models for the polyQ nucleus (Chen and Wolynes, 2017; Chen et al., 2016; Miettinen et al., 2014; T. T. M. Phan and Schmit, 2020), the AHA monomer contains an embryo of exquisitely ordered Q zipper structure necessary for the entropic bottleneck. We note that the AHA conformer is one of a small number of configurations for a minimal tertiary motif known as a beta-arcade, which was previously proposed to serve as an amyloid nucleus based on its favorable energetics and characteristic occurrence in amyloid fibrils (Kajava et al., 2010).

While our model may be impossible to confirm unequivocally -- as nuclei cannot be observed directly -- it nevertheless makes testable predictions. For example, it suggests that individual protofilaments can be as narrow as two or three zippered sheets -- approximately 1.6 or 2.4 nm, respectively. It also predicts spatial and temporal heterogeneities within polyQ aggregates, as a direct consequence of structural evolution, from short Q zippers with abundant connecting loops, to relatively long zippers with few loops. Specifically, at sufficiently high concentrations, spherulite-type aggregates can be expected with relatively disordered, soft interiors and ordered, hard exteriors. At sufficiently low concentrations, incipient protofilaments can be expected to grow in a unidirectional fashion. Some of these predictions have been borne out while this paper was in preparation. A cryoEM tomography study revealed a slab-like architecture of 51:Q amyloids in cells, with occasional restrictions to an apparent protofilament width of approximately 2 nm (Galaz-Montoya et al., 2021). Fluorescence microscopy studies of aging *C. elegans* neurons expressing Q:128, fused to the oligomerizing N-terminus of huntingtin and C-terminal CFP, revealed spherical aggregates with an apparently soft core and hard shell (Fisher et al., 2021).

That polyQ amyloid has a rate-limiting monomeric nucleus that is inhibited by oligomerization has important implications for disease etiology. First, it provides a rationale for why the age of onset for polyQ diseases appears not to depend on expression level nor copy number of the expanded polyQ allele. Second, it explains why the diseases instead depend on the length of the polyQ tract. Length thresholds for polyglutaminopathies tend to be in the 35–45 range (Matlahov and van der Wel, 2019). In agreement, we find that intramolecular nucleation occurs only for polyQ exceeding approximately 36 residues, corresponding to the length required to form a minimal intramolecular Q zipper.

### Interplay of Q zipper structure and surface tension may drive the accumulation of pre-amyloid oligomers

Soluble oligomers accumulate during the aggregation of pathologic length polyQ and/or Htt in vitro (Auer et al., 2008; Hsieh et al., 2017; Levin et al., 2014; Liang et al., 2018; Li et al., 2010; Sil et al., 2018; Vitalis and Pappu, 2011; Yamaguchi et al., 2005; Zanjani et al., 2020), in cultured cells (Olshina et al., 2010; Takahashi et al., 2008), and in the brains of patients (Legleiter et al., 2010; Sathasivam et al., 2010), and are likely culprits of proteotoxicity (Kim et al., 2016; Leitman et al., 2013; Lu and Palacino, 2013; Matlahov and van der Wel, 2019; Takahashi et al., 2008; Wetzel, 2020).

We found that Q zipper nucleation precedes the accumulation of soluble oligomers, rather than the other way around. These appeared to increase in size and order with the expression level of the protein, implying an evolution of the Q zipper structure toward mature amyloid. This progression rationalizes why polyQ diseases are rate-limited by primary nucleation despite amyloids -- the ultimate product of nucleation -- having benign or even protective roles (Kim et al., 2016; Leitman et al., 2013; Lu and Palacino, 2013; Matlahov and van der Wel, 2019; Takahashi et al., 2008; Wetzel, 2020).

As lamination maintains a lower aspect ratio than lengthening, the preference for polyQ, in a manner dependent on contralateral Qs, to oligomerize rather than elongate at low concentrations is plausibly mediated by the surface tension inherent to polyQ’s self-solvating property (Crick et al., 2006; Kang et al., 2017; Walters and Murphy, 2009). We speculate that this is a consequence of the fact that the opposition of surface tension to anisotropic growth falls above a critical curvature of the surface (Williamson et al., 2010). Hence, by inhibiting growth in one dimension, the removal of contralateral Qs increases the aspect ratio of Q zipper propagation and accelerates its escape from the surface tension-dominated regime. Flanking domains are likely to influence this balance (Crick et al., 2013; Williamson et al., 2010), and this presents an intriguing avenue for further research.

### A specific role for phase separation

Much recent work has revealed a tendency of amyloid-forming proteins, including those with Q-rich low complexity sequences, to undergo liquid-liquid phase separation under permissive conditions (Babinchak and Surewicz, 2020; Camino et al., 2021). PolyQ itself lacks both hydrophobic and charged residues that would confer the requisite affinity for phase separation in cellular milieu, which is rife with solubilizing competing interactors (Boncella et al., 2020; Gao et al., 2018; Wang et al., 2018; Xing et al., 2018). The physiological concentration of huntingtin in the human brain, estimated to be low nanomolar (Macdonald et al., 2014), is likely too low for liquid-liquid phase separation in the presence of even one binding partner at its physiological concentration (Posey et al., 2018). Notwithstanding limited reports to the contrary (Fisher et al., 2021; Peskett et al., 2018; Wan et al., 2021) -- which we respectfully attribute to known effects of flanking domain interactions and partitioning into pre-existing protein compartments (Duennwald et al., 2006a, 2006b; Jiang et al., 2017) -- our findings corroborate prior demonstrations that polyQ does not phase separate prior to amyloid formation, whether expressed in human neuronal cells (Colby et al., 2006; Kakkar et al., 2016), *C. elegans* body wall muscle cells (Sinnige et al., 2021), or [*pin*^-^] yeast cells (Duennwald et al., 2006a, 2006b; Jiang et al., 2017).

Counter to the prevailing paradigm, we found that oligomerization of naive polypeptides, whether from high concentration or fusion to a self-associating moiety, actually decelerated the rate-limiting step in amyloid formation -- nucleation of an incipient Q zipper. This likely relates to the fact that polyQ solvates itself better than water and consequently is already condensed intramolecularly (Crick et al., 2006; Walters and Murphy, 2009). That coalescence of those monomers into larger globules inhibited nucleation suggests that surface tension amplifies conformational fluctuations or even biases the conformational ensemble toward Q zippers. Prior molecular simulation studies reveal evidence of such effects. For example, polyQ polypeptides relax to a more expanded state within globules, with fewer reversals of chain direction, as a consequence of their becoming solvated by identical amides from other molecules rather than their own (Vitalis et al., 2009). Additionally, the side chains of Qs at the surface of polyQ globules are more dynamic and tend to orient toward the interior (Kang et al., 2017). Whether these changes correspond to backbone and side chain configurations that are more permissive to Q zipper formation remain to be determined.

We did find, however, that once an incipient Q zipper formed, phase separation strongly promoted its maturation to amyloid. This is consistent with lamination producing a multi-subunit templating surface; with the fact that increasing concentrations stabilize short beta strand interactions (T. M. Phan and Schmit, 2020); and with observations that polyQ amyloid elongation can occur without full consolidation to the fibril structure (Walters et al., 2012). To the extent that amyloid-associated soluble polyQ species drive disease, condensation would be more likely to ameliorate than exacerbate proteotoxicity.

### Concluding remarks

The etiology of polyQ pathology has been elusive. Tedious decades-long efforts by many labs have revealed precise measurements of polyQ aggregation kinetics, atomistic details of the amyloid structure, a catalogue of proteotoxic candidates, and snapshots of the conformational preferences of disordered polyQ. What those efforts notably have not led to, are treatments. Here, we synthesized those insights to recognize that pathogenesis likely begins with a very specific, and exceptionally rare, conformational fluctuation, and that our relative ignorance of that event may be blocking progress against polyQ diseases. We therefore set out to characterize the polyQ amyloid nucleus. Deploying a technique we recently developed to do just that, we arrived at a model for the single most important molecular species governing polyQ pathogenicity in vivo. We found that the model can explain key aspects of disease, such as length thresholds, kinetics of progression, and involvement of soluble amyloid-like species.

## Acknowledgments

We thank Viet Man and Jonathon Russell for assistance with molecular simulations; Jeremy Schmit, Ammon Posey, Rohit Pappu, Wei-feng Xue, and Patrick van der Wel for helpful discussions; Alejandro Rodriguez Gama for assistance with figure preparation; and Megan Halfmann for generous ancillary support. We thank Alexander Alexandrov for plasmids encoding Q_4_X repeats, early results from which stimulated this project’s inception. This work was funded by the National Institute Of General Medical Sciences of the National Institutes of Health under Award Number R01GM130927 (to RH) and the Stowers Institute for Medical Research. A portion of this work was done to fulfill, in part, requirements for a PhD thesis research for T.S.K. as a student registered with the Open University, UK at the Stowers Institute for Medical Research Graduate School, USA. Original data underlying this manuscript can be accessed from the Stowers Original Data Repository at http://www.stowers.org/research/publications/libpb-1494

## Materials and Methods

### Plasmid and Strain Construction

ORFs were codon optimized for expression in *S. cerevisiae,* synthesized, and cloned into vector V08 (Khan et al., 2018) by Genscript (Piscataway, NJ, USA).

Plasmids expressing fusions to MBP were created by inserting yeast-optimized MBP-SUMO to the N-terminus of the protein of interest. Plasmids expressing fusions to μNS were created by insertion of yeast-optimized μNS to the C-terminus of mEos3.1. See Table S1 for all full-length protein sequences.

Isogenic yeast strains rhy1713 ([*PIN^+^*]) and rhy1852 ([*pin*^-^]) were previously described (Khan et al., 2018). Yeast strain rhy2068 was created by homology-directed integration at the *ho* locus of rhy1713, a construct expressing *K. lactis URA3* from a tTA-tetO7 promoter (Bellí et al., 1998).

Yeast strains rhy2956 (MBP-32:Q_3_S), rhy2957 (MBP-60:Q_3_S), and rhy2958 (MBP-60:Q) were created by homologous recombination-mediated replacement of the *K. lactis URA3* ORF in rhy2068 with a PCR product of the coding sequence (minus mEos3.1) from plasmids rhx3735, rhx3736, and rhx3788, respectively.

### Yeast Preparation for DAmFRET, SDD-AGE and FACS

The yeast strains were transformed using a standard lithium acetate protocol with plasmids encoding the sequence to be tested as a fusion to the indicated tags (Table S1) under the control of the *GAL1* promoter.

Individual colonies were picked and incubated in 200μL of a standard synthetic media containing 2% dextrose (SD -ura) overnight while shaking on a Heidolph Titramax-1000 at 1000rpm at 30°C.

Following overnight growth, cells were washed twice with sterile water and resuspended in a synthetic induction media containing 2% galactose (SGal -ura). Cells were induced for 16 hours while shaking before being resuspended in fresh 2% SGal -ura for 4 hours to reduce autofluorescence. For DAmFRET, 75 μL of cells were then transferred to a 384 well plate for analysis on the cytometer.

### DAmFRET Cytometric Assay

Following 16 hours of total induction, cells were re-arrayed from a 96 well plate to a 384 well plate and photoconverted while shaking at 800rpm for 25 minutes using an OmniCure S1000 fitted with a 320-500 nm (violet) filter and a beam collimator (Exfo), positioned 45 cm above the plate. Cells were then transferred to a.Bio-Rad ZE5 cell analyzer.

For screening flow cytometry on the ZE5, plates were run in high-throughput mode (10 uL/well, flow speed 1.8) on the Bio-Rad ZE5 with Propel automation. Donor and FRET signals were collected from a 488 nm laser set to 100 mW, with voltages 351 and 370 respectively, into 525/35 and 593/52 filters.

Acceptor signal was collected from a 561 nm laser set to 50 mW with voltage at 525, into a 589/15 filter. Autofluorescence was collected from a 405 nm laser at 100 mW and voltage 450, into a 460/22 filter.

Compensation was performed manually, collecting files for non-photoconverted mEos3.1 for pure donor fluorescence and dsRed2 to represent acceptor signal as it has a similar spectrum to the red form of mEos3.1. FRET is compensated only in the direction of donor and acceptor fluorescence out of FRET channels, as there is not a pure FRET control.

Imaging flow cytometry was conducted as in previous work (Khan et. al 2018).

### DAmFRET Automated Analysis

FCS 3.1 files resulting from assay were gated using an automated R-script running in flowCore. Prior to gating, the forward scatter (FS00.A, FS00.W, FS00.H), side scatter (SS02.A), donor fluorescence (FL03.A) and autofluorescence (FL17.A) channels were transformed using a logicle transform in R. Gating was then done using flowCore by sequentially gating for cells using FS00.A vs SS02.A then selecting for single cells using FS00.H vs FS00.W and finally selecting for expressing cells using FL03.A vs FL17.A.

Gating for cells was done using a rectangular gate with values of Xmin = 2.7, Xmax = 4.8, Ymin = 2.7, Ymax = 4.7). Gating for single cells was done using a rectangular gate with values of Xmin = 4.45, Xmax = 4.58, Ymin = 2.5, Ymax = 4.4. Gating of expressing cells was done using a polygon gate with x/y vertices of (1,0.1), (1.8, 2), (5, 2), (5,0.1). Cells falling within all of these gates were then exported as FCS3.0 files for further analysis.

The FCS files resulting from the autonomous gating in Step 1 were then utilized for the JAVA-based quantification of a curve similar to the analysis found in the original assay (Khan et al 2018). Specifically:

The quantification procedure first divides a defined negative control DAmFRET histogram into 64 logarithmically spaced bins across a pre-determined range large enough to accommodate all potential data sets. Then upper gate values were determined for each bin as the 99^th^ percentile of the DamFRET distribution in that bin. For bins at very low and very high acceptor intensities, there are not enough cells to accurately calculate this gate value. As a result, for bins above the 99 ^th^ acceptor percentile and bins below 2 million acceptor intensity units, the upper gate value was set to the value of the nearest valid bin. The upper gate profile was then smoothed by boxcar smoothing with a width of 5 bins and shifted upwards by 0.028 DAmFRET units to ensure that the negative control signal lies completely within the negative FRET gate. The lower gate values for all bins were set equal to −0.2 DamFRET units. For all samples, then, cells falling above this negative FRET gate can be said to contain assembled (FRET-positive) protein. A metric reporting the gross percentage of the expressing cells containing assembled proteins is therefore reported as fgate which is a unitless statistic between 0 and 1.

This gate is then applied to all DAmFRET plots to define cells containing proteins that are either positive (self-assembled) or negative (monomeric). In each of the 64 gates, the fraction of cells in the assembled population were plotted as a ratio to total cells in the gate.

These values were then fit to a Weibull function from which the statistics EC50, δ and their respective errors (reported as the square of the residuals) were extracted (Khan et al. 2018). Mean δ and EC50 values are presented as the mean of four biological replicates with errors being represented by the standard deviation between replicates. We verified for all samples where conclusions are drawn from δ and/or EC50 that the Weibull fit closely approximated the raw DAmFRET plot.

In addition to these quantitations, we also utilized the mean raw AmFRET in each logarithmic window to detect even small changes in FRET between similar peptides agnostic to whether the assemblies were continuous or discontinuous.

### Microscopy

Cells were imaged using a CSU-W1 spinning disc Ti2 microscope (Nikon) and visualized through a 100x Plan Apochromat objective (NA 1.45). mEos3.1 was excited at 488 nm and emission was collected for 50 ms per frame through a ET525/36M bandpass filter on to a Flash 4 camera (Hamamatsu). Full z stacks of all cells were acquired over ~15 μm total distance with z spacing of 0.2 μm. Transmitted light was collected at the middle of the z stack for reference. To quantify the total intensity of each cell, the z stacks were processed using Fiji (https://imagej.net/software/fiji/). Images were first converted to 32-bit and sum projected in Z. Regions of interest (ROIs) were hand drawn around each cell using the ellipse tool in Fiji on the transmitted light image. These ROIs were then used to measure the area, mean, standard deviation, and integrated density of each cell on the fluorescence channel. Each cell was then classified as being diffuse or punctate by calculating the coefficient of variance (CV, standard deviation divided by the square root of the mean intensity) of the fluorescence. Cells were only considered punctate if their CV was greater than 30. Cells from each category that had equivalent integrated densities were directly compared and the images were plotted on the same intensity scale. The volume of each cell was calculated by fitting the transmitted light hand drawn ROI to an ellipse. The cell was then assumed to be a symmetric ellipsoid with the parameters of the fit ellipse from Fiji. The volume was calculated by 4/3*pi*major*minor*minor, where major and minor are the major and minor axes of the Fiji ellipse fit to the hand drawn ellipse ROI. The volumes reported are the volumes of the 3D ellipsoids. Concentrations were calculated by dividing the integrated densities by the calculated volume in μm^3^, yielding units of fluorescence per μm^3^.

### SDD-AGE

Semi-denaturing detergent agarose gel electrophoresis (SDD-AGE) was done as in (Khan et al., 2018). The gel was imaged directly using a GE Typhoon Imaging System using a 488 laser and 525(40) BP filter. Images were then loaded into ImageJ for contrast adjustment. Images were gaussian blurred with a radius of 1 and then background subtracted with a 200 pixel rolling ball. Representative samples were then cropped from the original image for emphasis. Dotted or solid lines denote where different regions of the same gel were aligned for comparison.

### Bioinformatic Analysis

Comparative analyses were performed using default settings at the listed, public web servers as were available on April 1st, 2021. See Table S2 for further details.

### Molecular Simulations

The simulations were carried out using the AMBER 20 package (Case et al., 2020) with ff14SB force field (Maier et al., 2015) and explicit TIP3P water model (Price and Brooks, 2004). The aggregates were placed in a cubic solvation box. In each aggregate, the minimum distance from the aggregate to the box boundary was set to be 12 nm to avoid self-interactions. An 8 Å cutoff was applied for the nonbonded interactions, and the particle mesh Ewald (PME) method (Essmann et al., 1995) was used to calculate the electrostatic interaction with cubic-spline interpolation and a grid spacing of approximately 1 Å. Once the box was set up, we performed a structural optimization with aggregates fixed, and on a second step allowed them to move, followed by a graduate heating procedure (NVT) with individual steps of 200 ps from 0 K to 300 K. Finally, the production runs were carried out using NPT Langevin dynamics with constant pressure of 1 atm at 300 K.

## Supplemental Files

**Table S1**. **List of plasmids and sequences.**

**Table S2. Amyloid predictor output.**

**Table S3. Weibull fit statistics for L:Q_3_S variants**.

**Figure S1. Anatomy of an amyloid nucleation barrier.**

A. Reaction coordinate diagram schematizing the energy barrier for forming a hypothetical homogeneous amyloid nucleus (blue circle with red square). Because amyloid involves a transition from disordered monomer (left) to ordered multimer (right), the nucleation barrier (purple) is a combination of high energy fluctuations in both density (blue) and conformation (red).

B. Increased local concentrations of the protein make the critical fluctuation in density, specifically, more favorable (gray arrows indicate reduction in ΔG), increasing the frequency of nucleation while making it more dependent on conformational fluctuations.

C. Conformational templates, such as amyloids of a different protein ([*PIN^+^*]), make the critical fluctuations in conformation, specifically, more favorable, increasing the frequency of nucleation while making it more dependent on density fluctuations.

**Figure S2.**

A. DAmFRET plots of [*pin*^-^] and [*PIN*^+^] cells expressing sequence variants with greater than 50% N, showing negligible nucleation in the absence of a conformational template. Shown are representative plots from the same experiment.

B. DAmFRET plots of [*pin*^-^] and [*PIN*^+^] cells expressing polypeptides composed of tandem repeats of *a* Qs separated by an N for a total length of 60 residues. Shown are representative plots from the same experiment.The plots surrounded by a purple dashed box illustrate the striking sequence parity of de novo nucleation potential.

C. DAmFRET plots of [*pin*^-^] cells expressing Q_3_X, Q_4_X, and Q_5_X sequences where X = N, G, A, S, or H. H-containing repeat sequences are length 75; all others are length 60. Shown are representative plots from the same experiment.

D. DAmFRET plots of [*PIN^+^*] cells expressing Q_3_X, Q_4_X, and Q_5_X sequences where X = N, G, A, S, or H. H-containing repeat sequences are length 75; all others are length 60. Shown are representative plots from the same experiment

**Figure S3.**

A. Molecular simulations of model Q zippers formed by a pair of two-stranded antiparallel beta sheets, containing a single serine residue (QQQSQQQ) per strand. The structure is unstable when the S side chains face inward (top), but not when the S side chains face outward (bottom).

B. Simulations of model steric zippers formed by a pair of four-stranded parallel beta sheets, containing various numbers and orientations of non-Q residues. From top to bottom: one asparagine inside, one serine inside, multiple asparagines outside, multiple serines outside. The structures proved unstable in all cases, indicating that amyloid fibers formed by Q_(<8)_X repeat polypeptides are unlikely to have a parallel beta sheet arrangement.

C. Schematic demonstrating how a polar clasp (red dashed line) would preclude Q zipper formation.

D. Schema and frequencies of polar clasps occurring between two unilaterally adjacent Q side chains (left) or a unilaterally adjacent N and Q side chain (right) within a QQQNQQQ peptide, simulated either with (top) or without (bottom) the backbone restrained in a beta conformation. The bar graphs show that polar clasps between Q and N occur less frequently than between Q and Q, indicating that the mechanism of Q zipper destabilization by N side chains cannot be attributed to polar clasps.

E. Schema and lifetimes of H-bonds between exterior stacked (axially adjacent) Q side chains (top) or N and Q side chains (bottom) in the Q zipper simulated in Fig. 2D, showing no difference in stabilities between axial H-bonds between Q and Q/N side chains.

F. As a consequence of the N side chain’s interception of the opposing Q side chain’s H-bond, the Q is no longer anchored in the outstretched configuration and sterically interferes with the ordering of adjacent Qs. This effect propagates through the zipper, resulting in its dissolution.

G. As for N, the side chain of S is too short to H-bond with the opposing main chain. Unlike for N, however, the S side chain is also too short to intercept the opposing Q side chain’s H-bond, allowing the Q to H-bond (black arrow) the backbone amide adjacent to the S. Therefore, whereas Q zippers cannot accommodate internal N residues, they can accommodate sparse internal S residues.

**Figure S4.**

A. SDD-AGE of the indicated mEos3.1-tagged proteins expressed in both [*PIN*^+^] and [*pin*^-^] cells. Lysates were normalized by fluorescence to within 50% of each other prior to loading. Sample buffer was added to a final concentration of 2% either SDS (top) or sarkosyl (bottom). Sarkosyl is a milder detergent that allows for detection of amyloids that are denatured by SDS.

B. DAmFRET plots of [*pin*^-^] cells expressing unilateral contiguity variants of the 60:Q_3_N base sequence, showing that at least six unilaterally contiguous glutamines (see schematic) are required for *de novo* nucleation of single long Q zipper amyloid with low concentration-dependence.

C. DAmFRET plots of contralateral contiguity variants of the 60:Q_4_N base sequence, showing that at least six bilaterally contiguous Qs are required for robust amyloid formation.

**Figure S5.**

A. DAmFRET plots of 60:Q_5_N and 60:Q_8_N acquired using imaging flow cytometry, showing gates at high expression for both high- or low-AmFRET populations. Insets show from left to right the distribution of donor, FRET, and acceptor fluorescence, respectively, in representative cells from each gate.

B. DAmFRET plots of [*pin*^-^] cells expressing the indicated proteins with and without genetic fusion to μNS.

C. DAmFRET plots of [*PIN*^-^] cells expressing the indicated proteins with and without genetic fusion to μNS.

**Figure S6.**

A. DAmFRET plots of [*PIN*^+^] cells expressing ASC, with or without fusion to MBP. Right plot shows the fraction of cells FRET-positive and the corresponding Weibull fits for triplicates, with inset reporting statistics of the Weibull fits +/- error. MBP specifically increased the C_min_ and EC_50_ values.

B. DAmFRET plots of [*pin*^-^] and [*PIN*^+^] cells expressing the indicated sequences, with or without fusion to MBP.

C. DAmFRET plots of 60:Q_3_N (top) or 60:Q_7_N (bottom) in [*PIN^+^*] cells either lacking or expressing non-fluorescent MBP-32:Q_3_S or MBP-60:Q_3_S in trans. 60:Q_3_N nucleates at lower concentration specifically when seeded by MBP-60:Q_3_S. Black arrow shows the appearance of a mid-FRET population for 60:Q_7_N, indicative of single Q zipper amyloids, only when seeded by MBP-60Q_3_S. Dashed boxes correspond to the histograms in Fig. 6D.

**Figure S7.**

A. DAmFRET plots (left) and corresponding output from automated Weibull fitting (middle and right) of the indicated Q_3_S length variants expressed in [*PIN^+^*] cells. Note that the Weibull fits in this case assume monomeric nucleation, i.e. the C_min_ parameter was constrained to 0, in order to tighten the confidence intervals for the other two parameters. The region outlined in yellow in the middle plots designates the non-FRET population used to distinguish whether nucleation has (outside) or has not (inside) occurred in each cell. The black trace in the right plots shows the fraction of cells nucleated in different concentration windows, and the blue trace shows the corresponding Weibull fit.

B. DAmFRET plots of polyQ length variants in [*PIN^+^*] cells.

C. DAmFRET plots of polyQ length variants in [*pin*^-^] cells.

D. Spline fits of DAmFRET data for polyQ lengths 8-35 expressed in [*PIN^+^*] and [*pin*^-^] cells. Only at 35:Q does AmFRET significantly increase relative to 8:Q (p < 0.000001; t-test), and only in [*PIN^+^*] cells.

E. DAmFRET plots of μNS-fused polyQ length variants.

**Figure S8.** Minimal Q zipper conformers of polyQ compatible with our experimental data. Left columns contain a side view of the structures after equilibration, and a scheme of the topology of the beta strands as seen from the left end (cross, strand into the plane; dot, strand coming out of the plane).

Vertical dashed lines represent rows of hydrogen bonds within sheets (H-bonds involving side chains are not shown). Right columns show the time evolution of the structures, initial configurations are shown after energy minimization and equilibration. In all cases, the fibril axes are in the direction of the hydrogen bonds (vertical dashed lines).

A. One of the hypothetical folds consistent with all constraints identified for the polyQ amyloid nucleus. This fold differs from B in having the first and fourth strands unpaired and flanking either side of the central hairpin. The initial conformer in this simulation had 11 residues per strand. Note that the visualization package does not plot unpaired strands as ribbons.

B. Another hypothetical fold consistent with all constraints identified for the polyQ amyloid nucleus. This fold, dubbed AHA, differs from A in having the first and fourth strands paired and flanking only one side of the central hairpin. The initial conformer in this simulation had 8 residues per strand; the stable conformer (run for 1 □s) had 6 residues per strand.

C. Minimal monomeric subunit of single Q zipper amyloid. It comprises a beta hairpin and a flanking third extended strand. The initial conformer in this simulation had 11 residues per strand.

D. Axial homodimer of C, representing the smallest hypothetically possible Q zipper amyloid dimer.

## Notes

### Competing Interest Statement

The authors have declared no competing interest.

### Summary of Updates

This version of the manuscript has been revised to include additional citations and improve readability.

