## Supplemental Figures for "The polyglutamine amyloid nucleus in living cells is a monomer with competing dimensions of order"

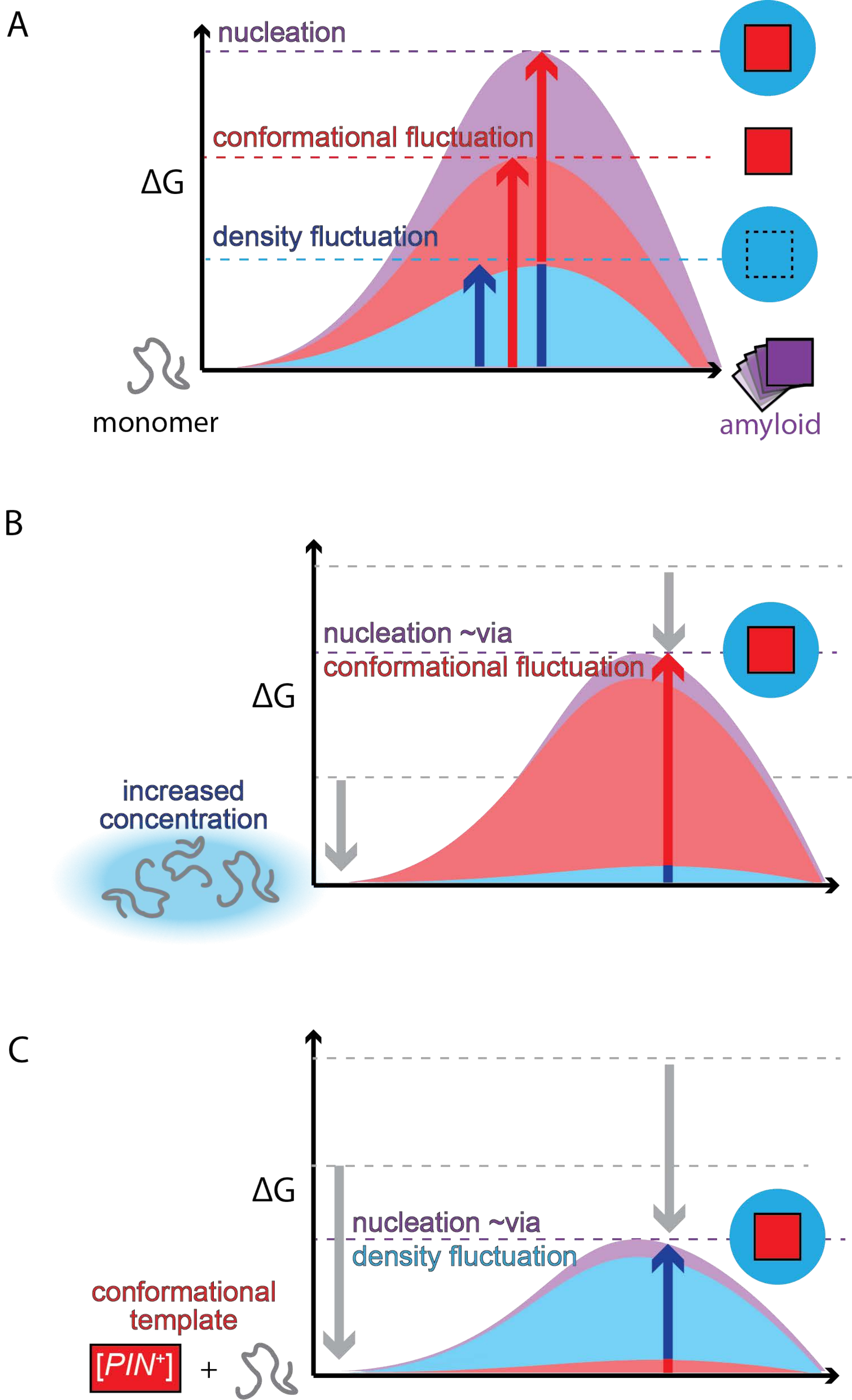

A
nucleation
conformational fluctuation
ΔG
density fluctuation
monomer
amyloid
B
nucleation ~via
conformational fluctuation
ΔG
increased
concentration
C
ΔG
nucleation ~via
density fluctuation
conformational
template
[PIN+]
+

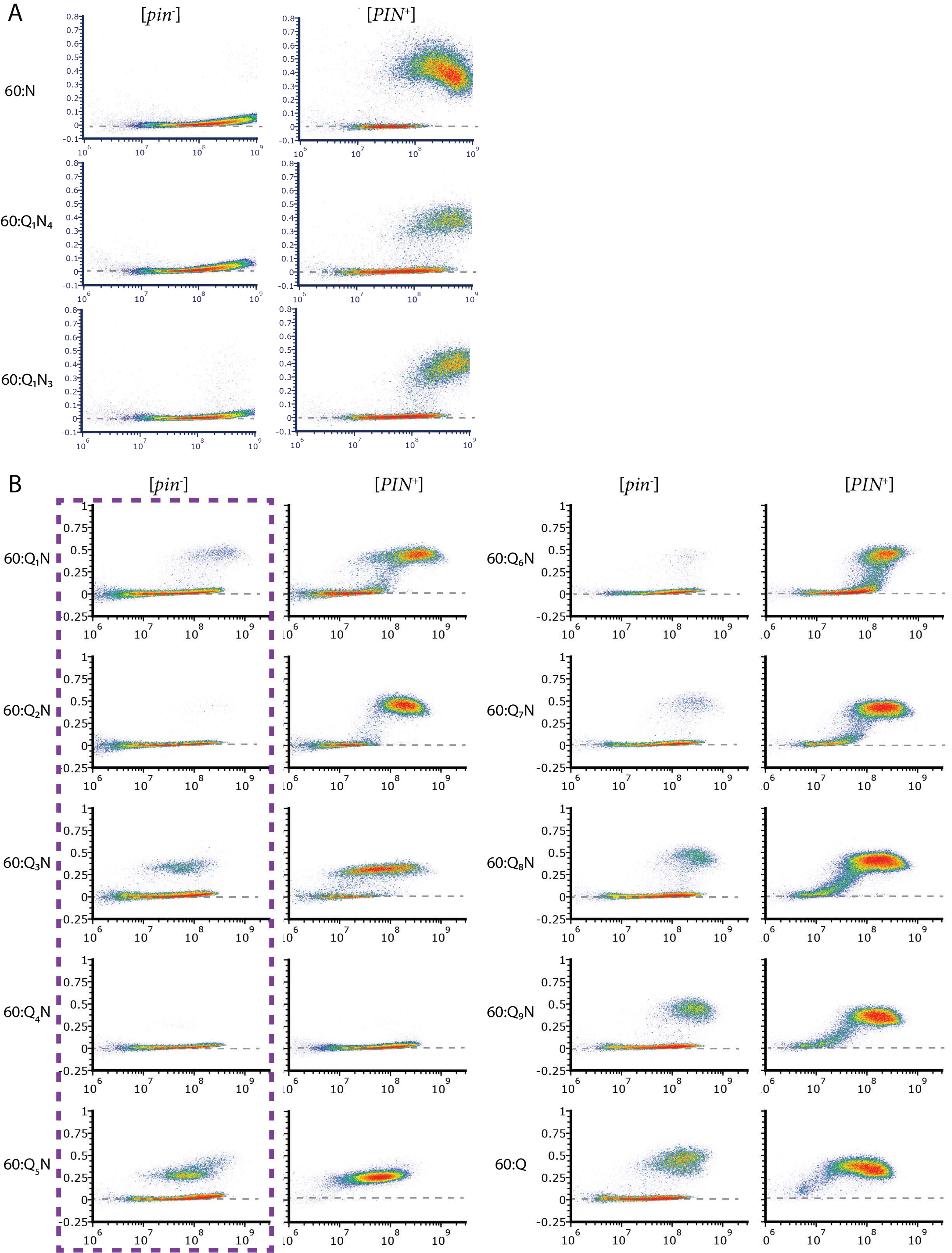

A
[pin-]
[PIN+]
60:N
60:Q1N4
60:Q1N3
B
[pin-]
[PIN+]
[pin-]
[PIN+]
60:Q1N
60:Q2N
60:Q3N
60:Q4N
60:Q5N
60:Q6N
60:Q7N
60:Q8N
60:Q9N
60:Q

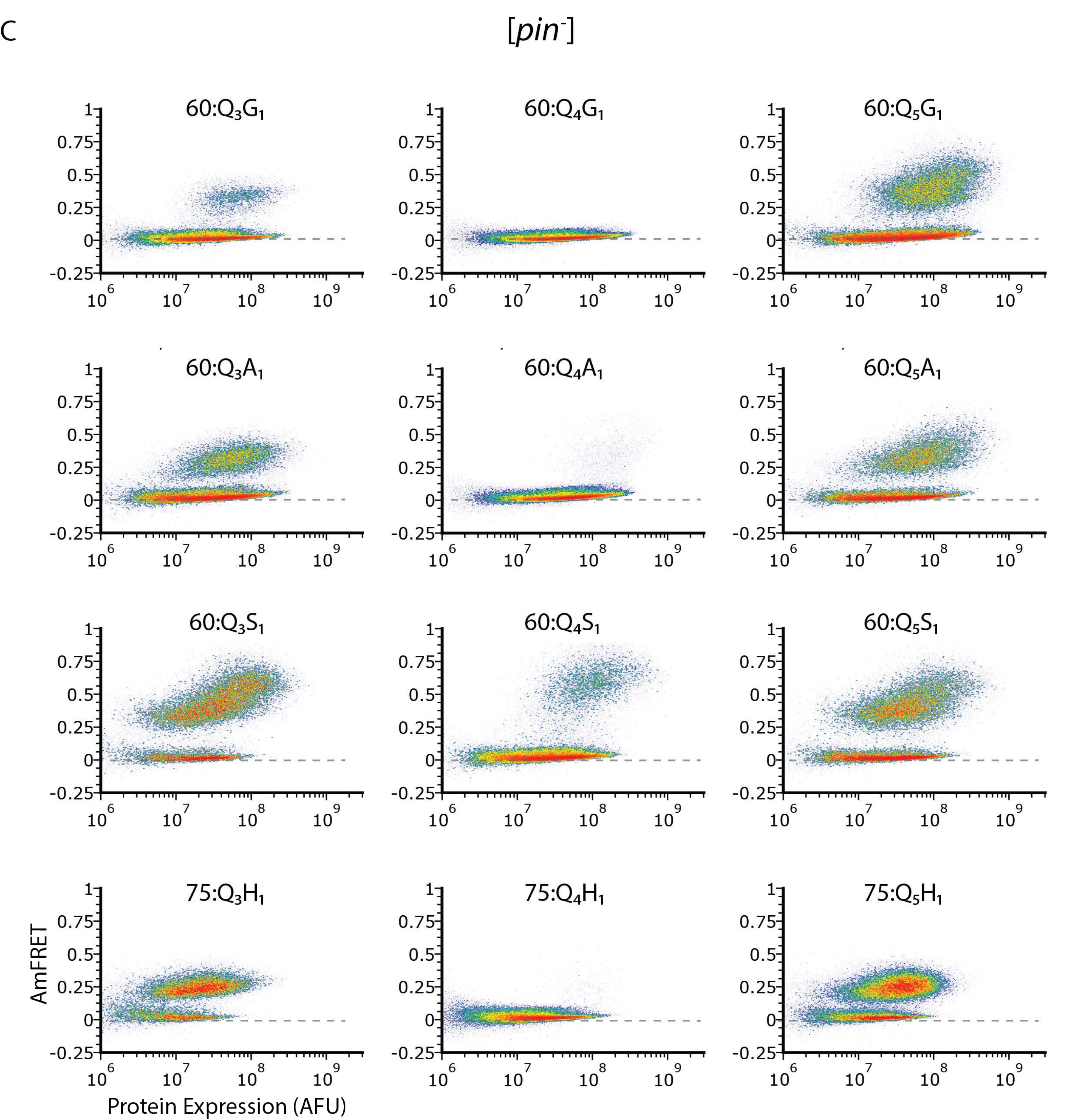
C
[pin-]
60:Q3G1
60:Q4G1
60:Q5G1
60:Q3A1
60:Q4A1
60:Q5A1
60:Q3S1
60:Q4S1
60:Q5S1
75:Q3H1
75:Q4H1
75:Q5H1
AmFRET
Protein Expression (AFU)
1
0.75
0.5
0.25
0
-0.25
10^6
10^7
10^8
10^9

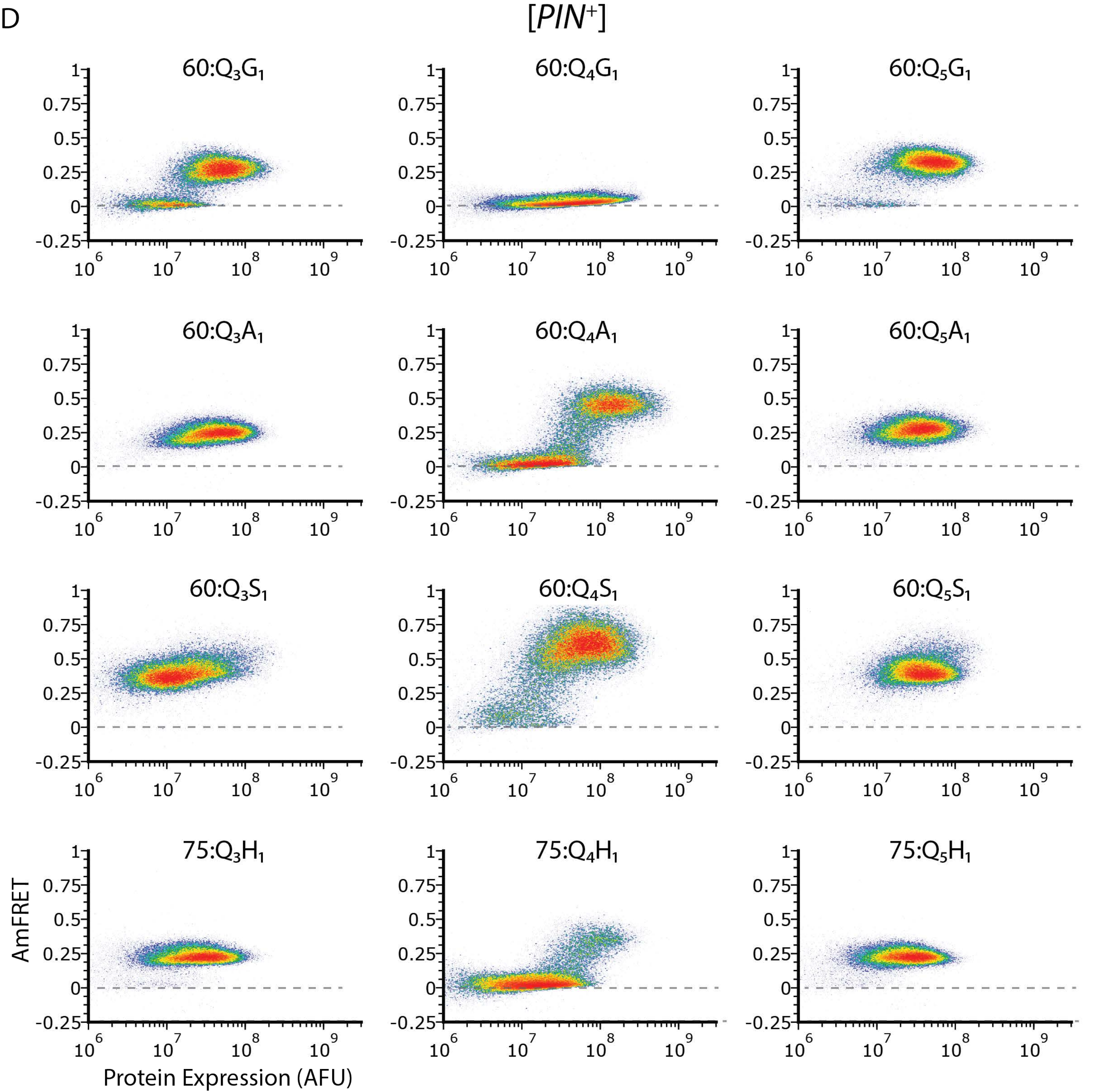
D
[PIN+]
60:Q3G1
60:Q4G1
60:Q5G1
60:Q3A1
60:Q4A1
60:Q5A1
60:Q3S1
60:Q4S1
60:Q5S1
75:Q3H1
75:Q4H1
75:Q5H1
AmFRET
Protein Expression (AFU)

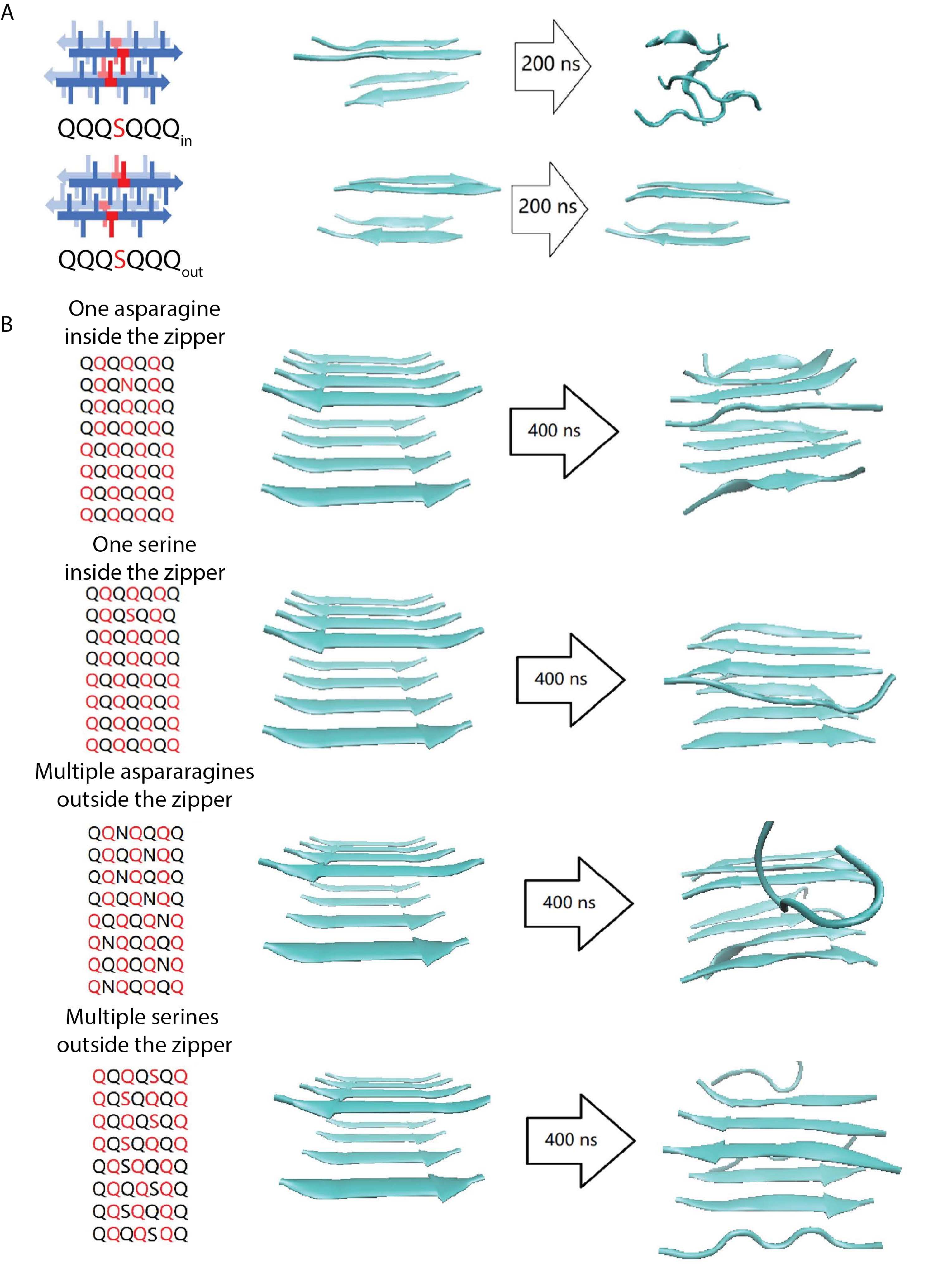

A
QQQSQQQin
200 ns
QQQSQQQout
200 ns
B
One asparagine
inside the zipper
QQQQQQQ
QQQNQQQ
QQQQQQQ
QQQQQQQ
QQQQQQQ
QQQQQQQ
QQQQQQQ
QQQQQQQ
400 ns
One serine
inside the zipper
QQQQQQQ
QQQSQQQ
QQQQQQQ
QQQQQQQ
QQQQQQQ
QQQQQQQ
QQQQQQQ
QQQQQQQ
400 ns
Multiple aspararagines
outside the zipper
QQNQQQQ
QQQQNQQ
QQNQQQQ
QQQQNQQ
QQQQQNQ
QNQQQQQ
QQQQQNQ
QNQQQQQ
400 ns
Multiple serines
outside the zipper
QQQQSQQ
QQSQQQQ
QQQQSQQ
QQSQQQQ
QQSQQQQ
QQQQSQQ
QQSQQQQ
QQQQSQQ
400 ns

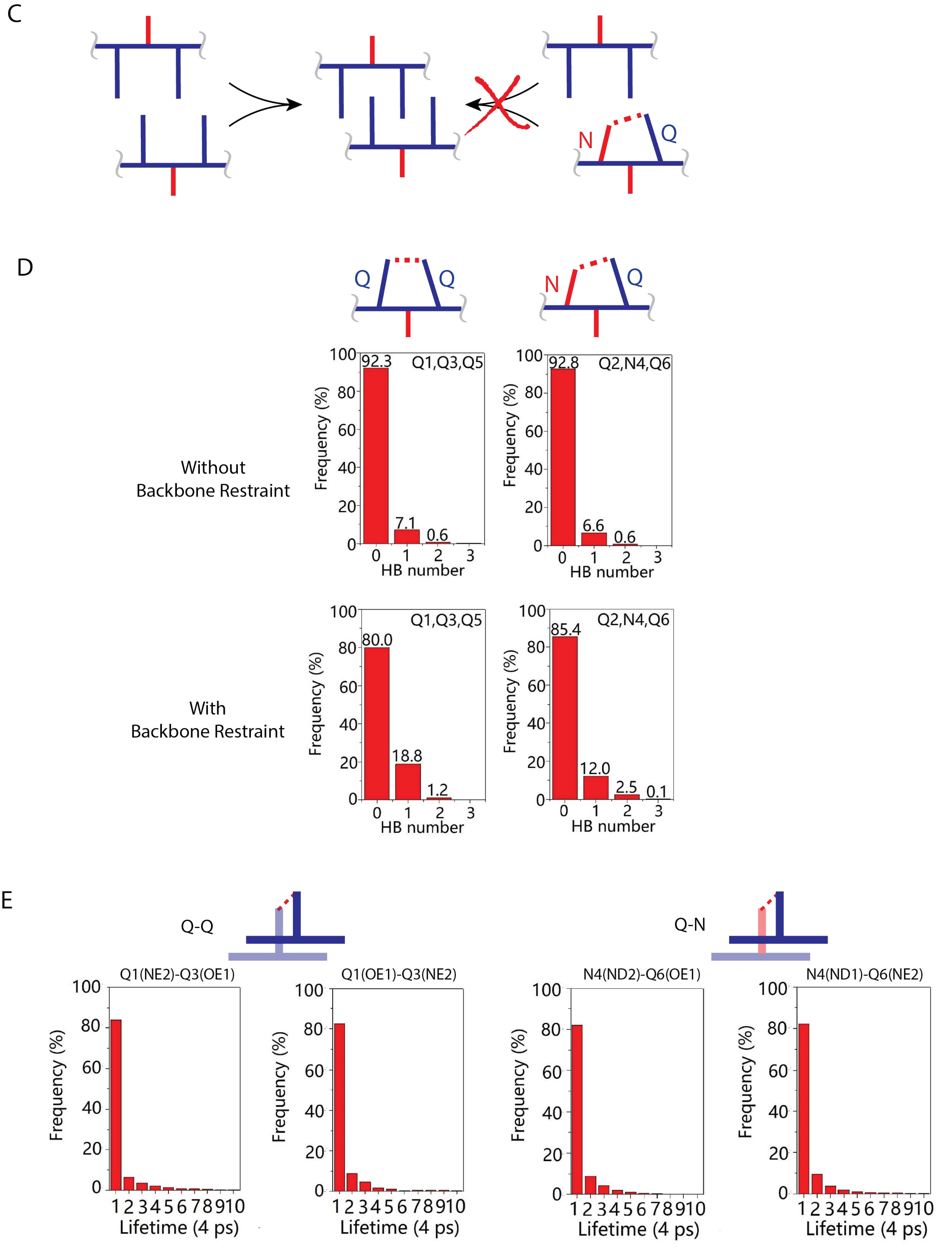
C
N
Q
D
Q
Q
N
Q
Without
Backbone Restraint
With
Backbone Restraint
Q1,Q3,Q5
Q2,N4,Q6
Frequency (%)
HB number
92.3
7.1
0.6
92.8
6.6
0.6
80.0
18.8
1.2
85.4
12.0
2.5
0.1
E
Q-Q
Q-N
Q1(NE2)-Q3(OE1)
Q1(OE1)-Q3(NE2)
N4(ND2)-Q6(OE1)
N4(ND1)-Q6(NE2)
Lifetime (4 ps)

F

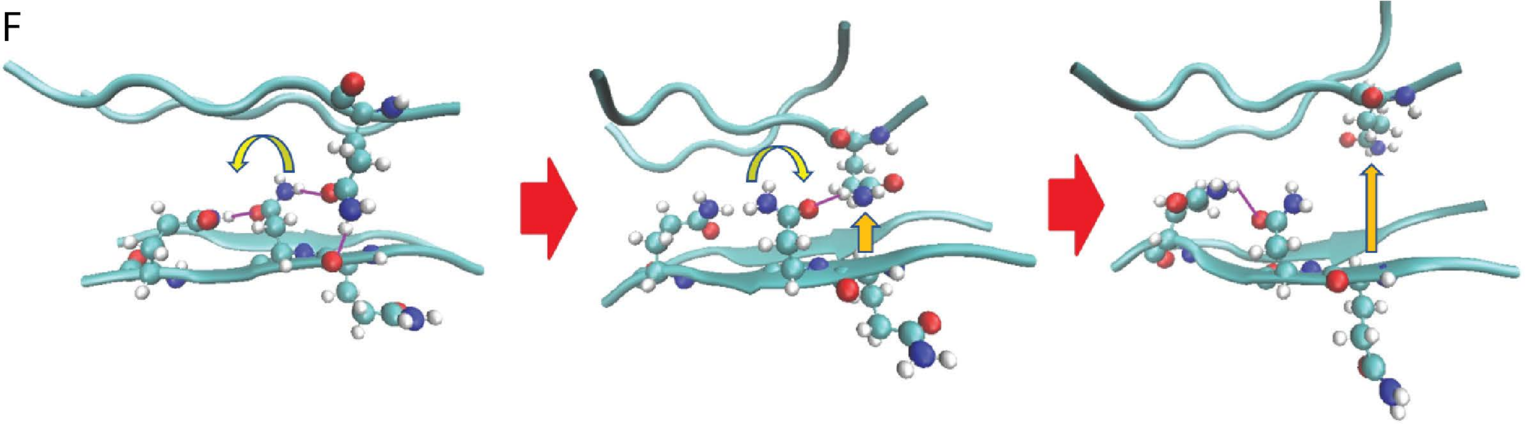

G

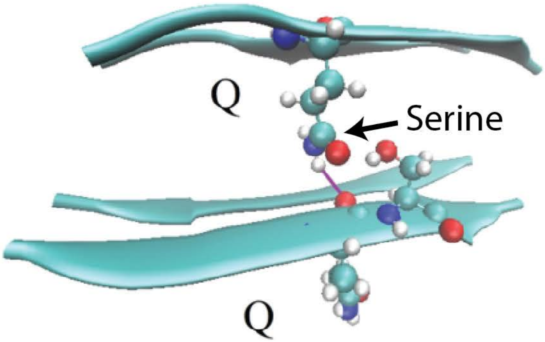

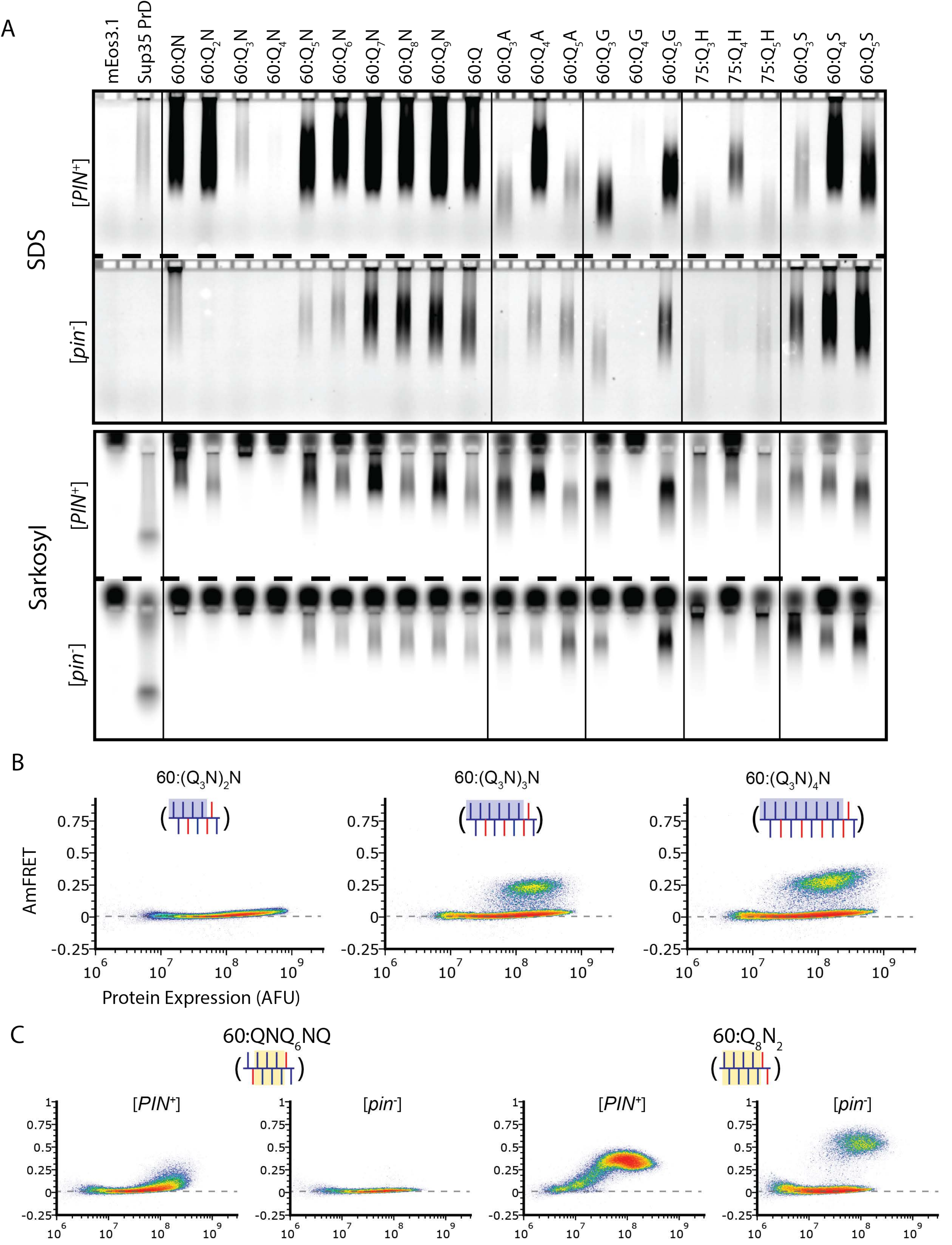
A
mEos3.1
Sup35 PrD
60:QN
60:Q2N
60:Q3N
60:Q4N
60:Q5N
60:Q6N
60:Q7N
60:Q8N
60:Q9N
60:Q
60:Q3A
60:Q4A
60:Q5A
60:Q3G
60:Q4G
60:Q5G
75:Q3H
75:Q4H
75:Q5H
60:Q3S
60:Q4S
60:Q5S
SDS
[PIN+]
[pin-]
Sarkosyl
[PIN+]
[pin-]
B
60:(Q3N)2N
60:(Q3N)3N
60:(Q3N)4N
AmFRET
Protein Expression (AFU)
C
60:QNQ6NQ
60:Q8N2
[PIN+]
[pin-]
[PIN+]
[pin-]

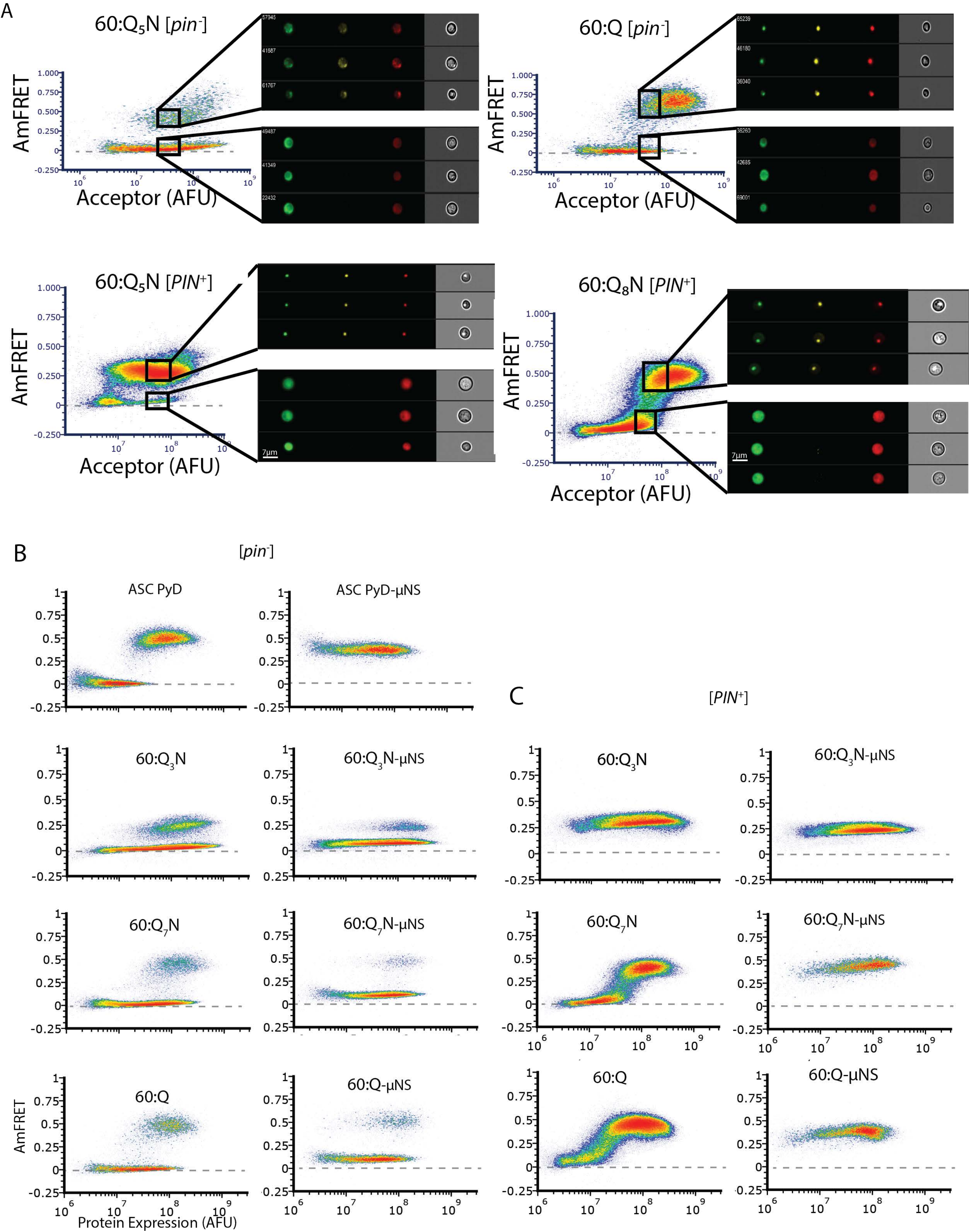

A
60:Q5N [pin-]
60:Q [pin-]
60:Q5N [PIN+]
60:Q8N [PIN+]
AmFRET
Acceptor (AFU)
7µm
B
[pin-]
ASC PyD
ASC PyD-µNS
60:Q3N
60:Q3N-µNS
60:Q7N
60:Q7N-µNS
60:Q
60:Q-µNS
AmFRET
Protein Expression (AFU)
C
[PIN+]
60:Q3N
60:Q3N-µNS
60:Q7N
60:Q7N-µNS
60:Q
60:Q-µNS

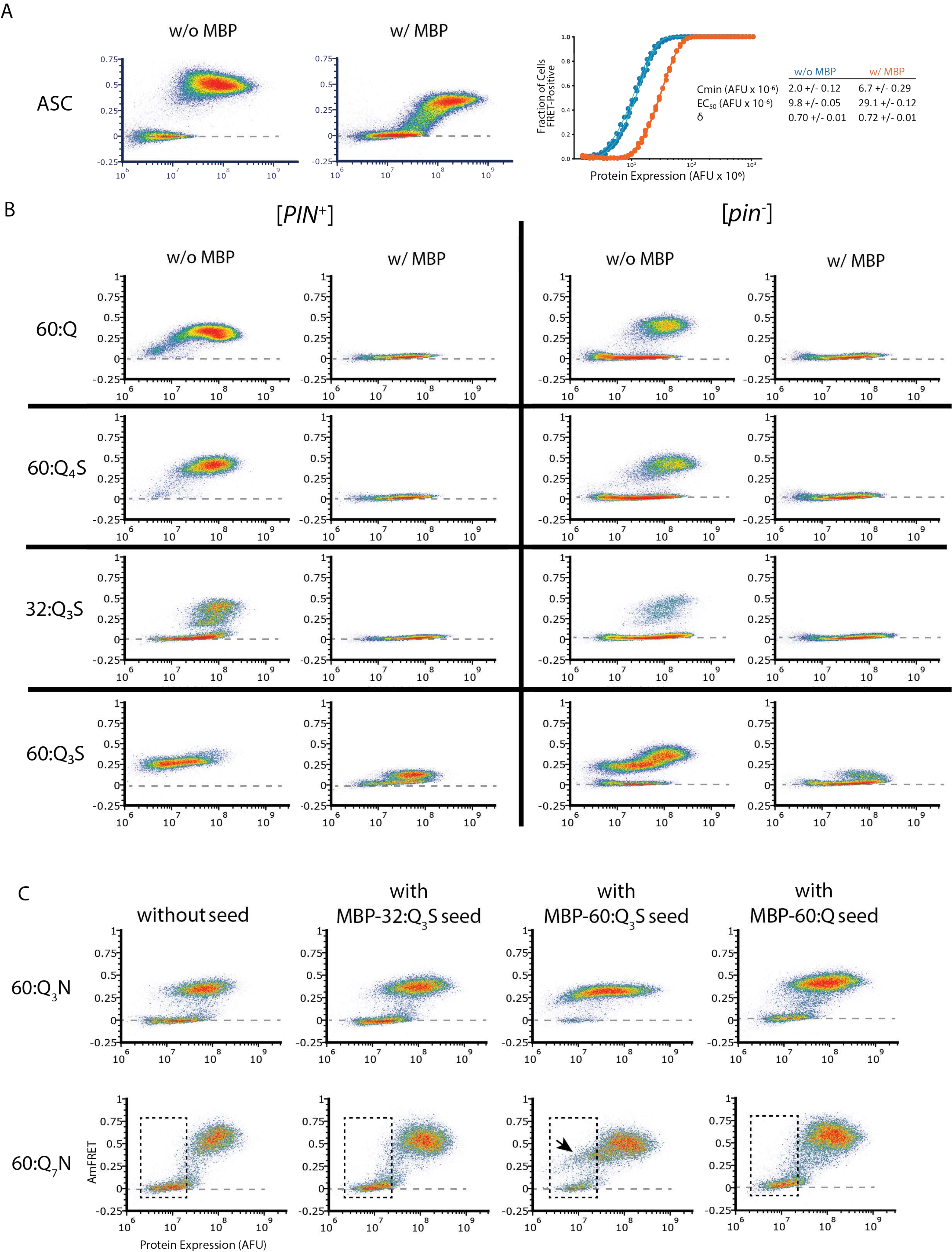

A
w/o MBP
w/ MBP
ASC
Fraction of Cells FRET-Positive
Protein Expression (AFU x 10^6)
w/o MBP
w/ MBP
Cmin (AFU x 10^-6) 2.0 +/- 0.12 6.7 +/- 0.29
EC50 (AFU x 10^-6) 9.8 +/- 0.05 29.1 +/- 0.12
δ 0.70 +/- 0.01 0.72 +/- 0.01
B
[PIN+]
[pin-]
w/o MBP
w/ MBP
60:Q
60:Q4S
32:Q3S
60:Q3S
C
without seed
with MBP-32:Q3S seed
with MBP-60:Q3S seed
with MBP-60:Q seed
60:Q3N
60:Q7N
AmFRET
Protein Expression (AFU)

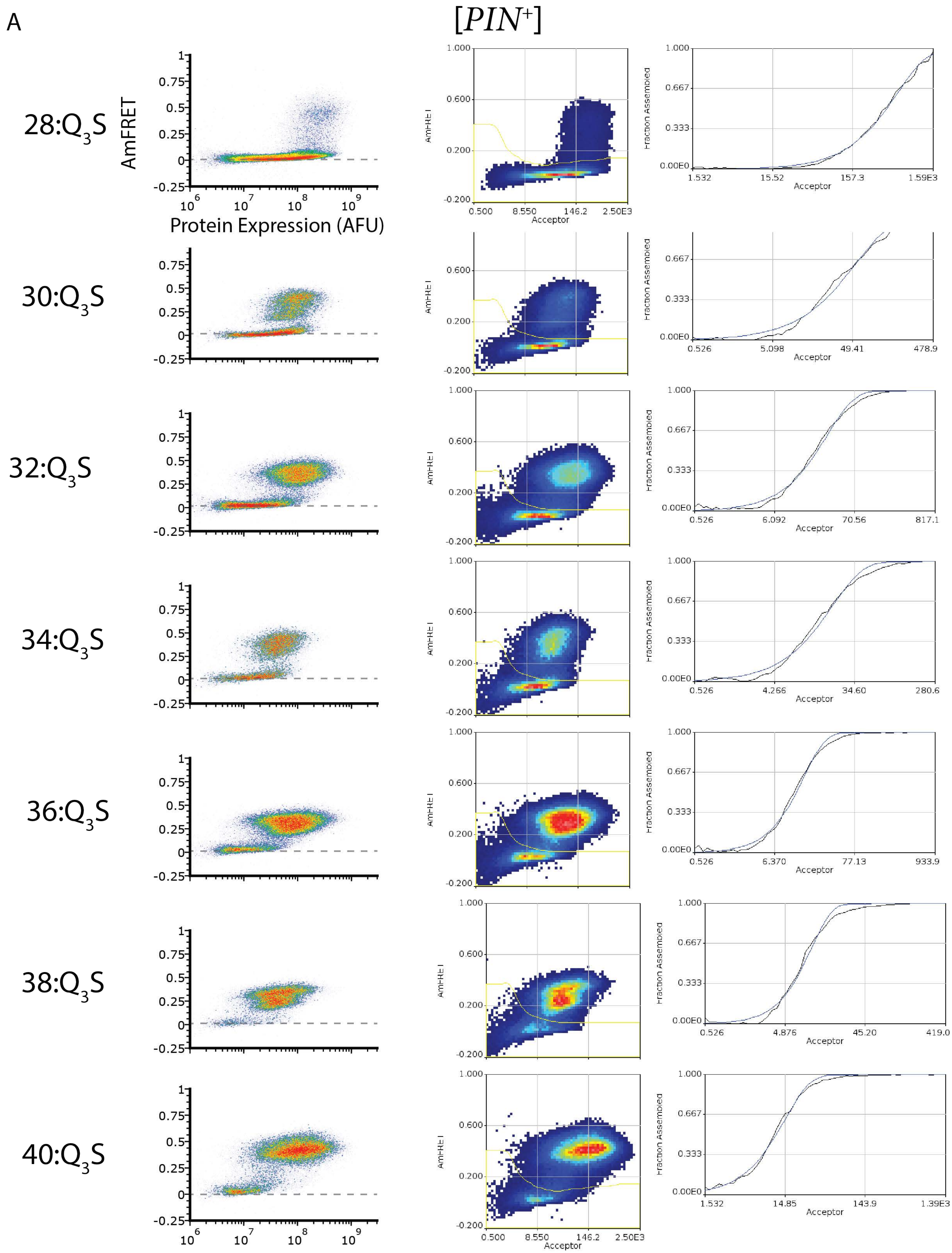

A
[PIN+]
28:Q3S
30:Q3S
32:Q3S
34:Q3S
36:Q3S
38:Q3S
40:Q3S
AmFRET
Protein Expression (AFU)
Acceptor
Fraction Assembled
1.532
15.52
157.3
1.59E3
0.526
5.098
49.41
478.9
6.092
70.56
817.1
4.266
34.60
280.6
6.370
77.13
933.9
4.876
45.20
419.0
14.85
143.9
1.39E3
0.500
8.550
146.2
2.50E3

B

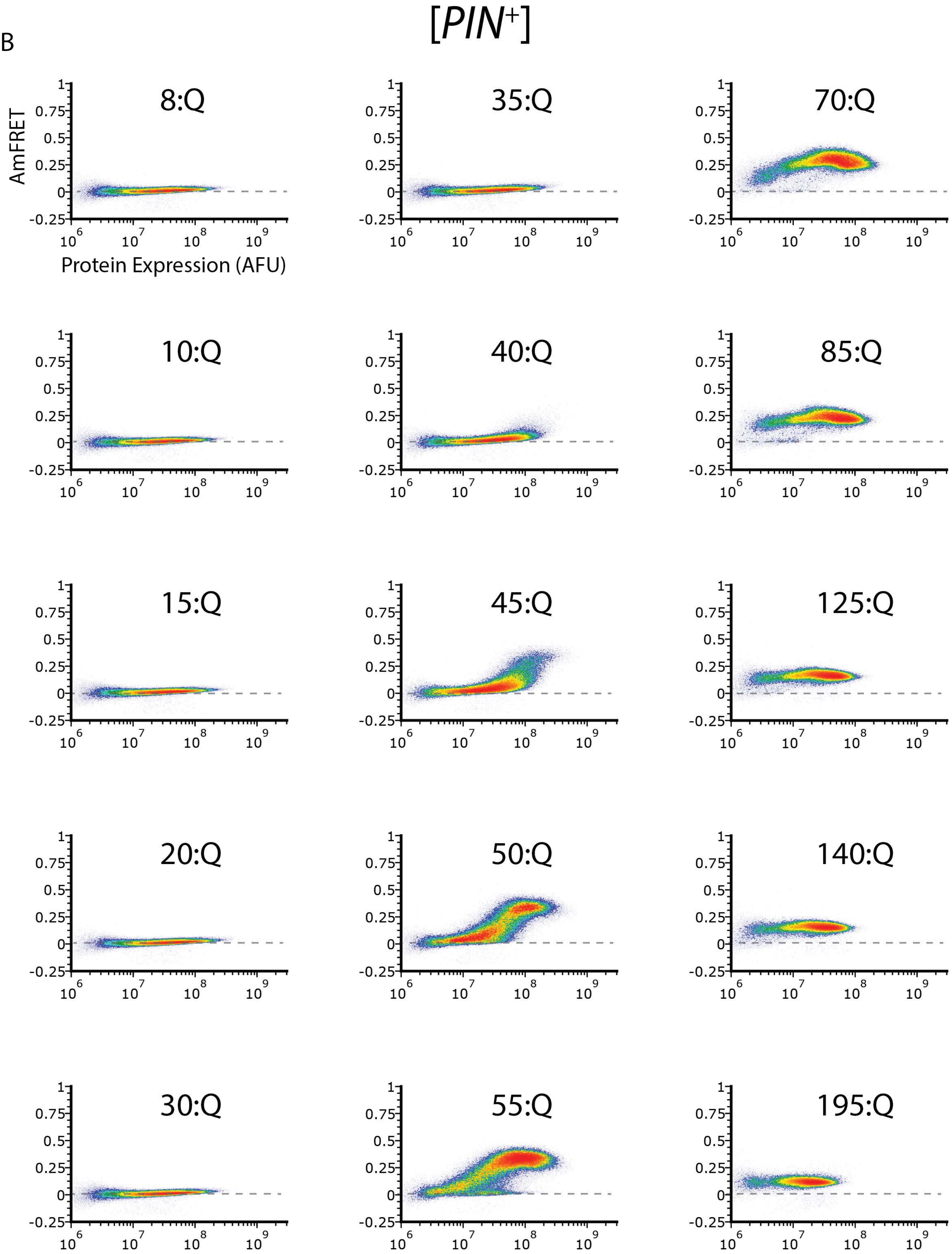

[PIN+]
AmFRET
Protein Expression (AFU)
8:Q
35:Q
70:Q
10:Q
40:Q
85:Q
15:Q
45:Q
125:Q
20:Q
50:Q
140:Q
30:Q
55:Q
195:Q

C

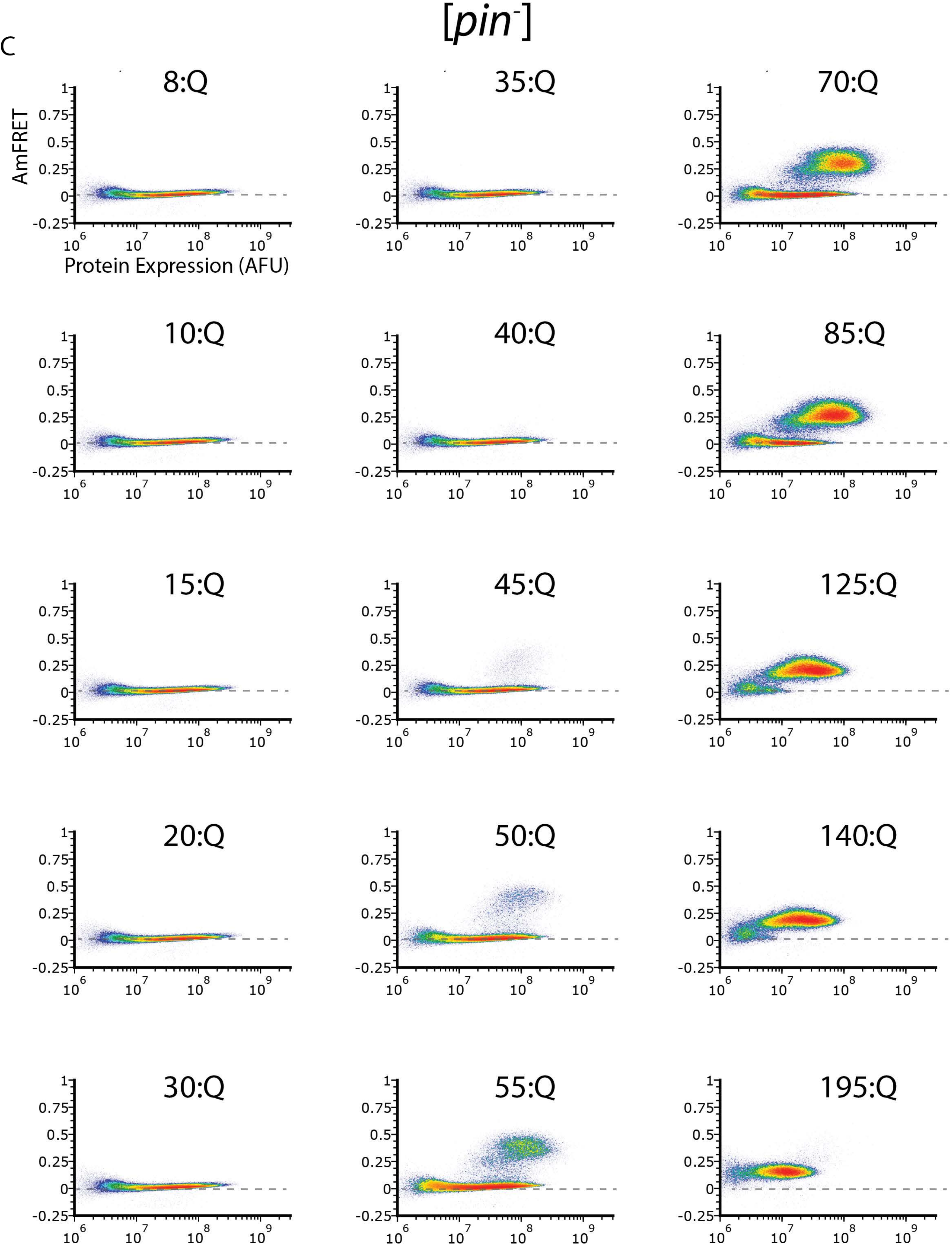

[pin-]
8:Q
35:Q
70:Q
10:Q
40:Q
85:Q
15:Q
45:Q
125:Q
20:Q
50:Q
140:Q
30:Q
55:Q
195:Q
AmFRET
Protein Expression (AFU)

D

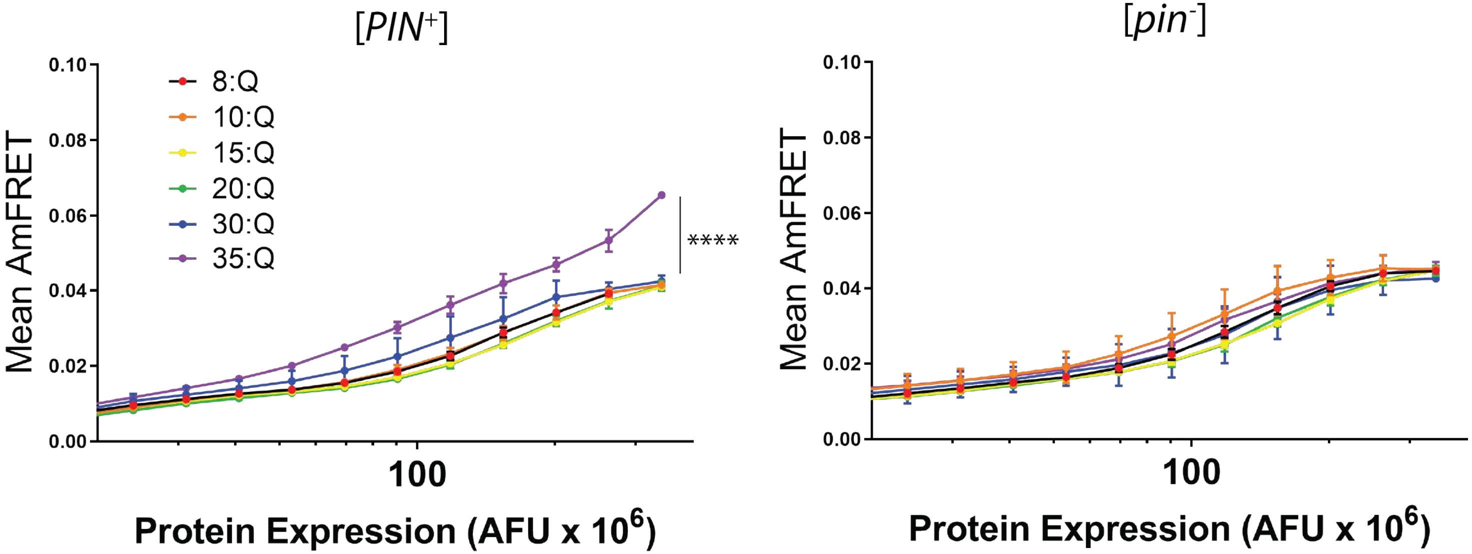

[PIN+]
[pin-]
Mean AmFRET
0.10
0.08
0.06
0.04
0.02
0.00
8:Q
10:Q
15:Q
20:Q
30:Q
35:Q
****
100
Protein Expression (AFU x 10^6)


E

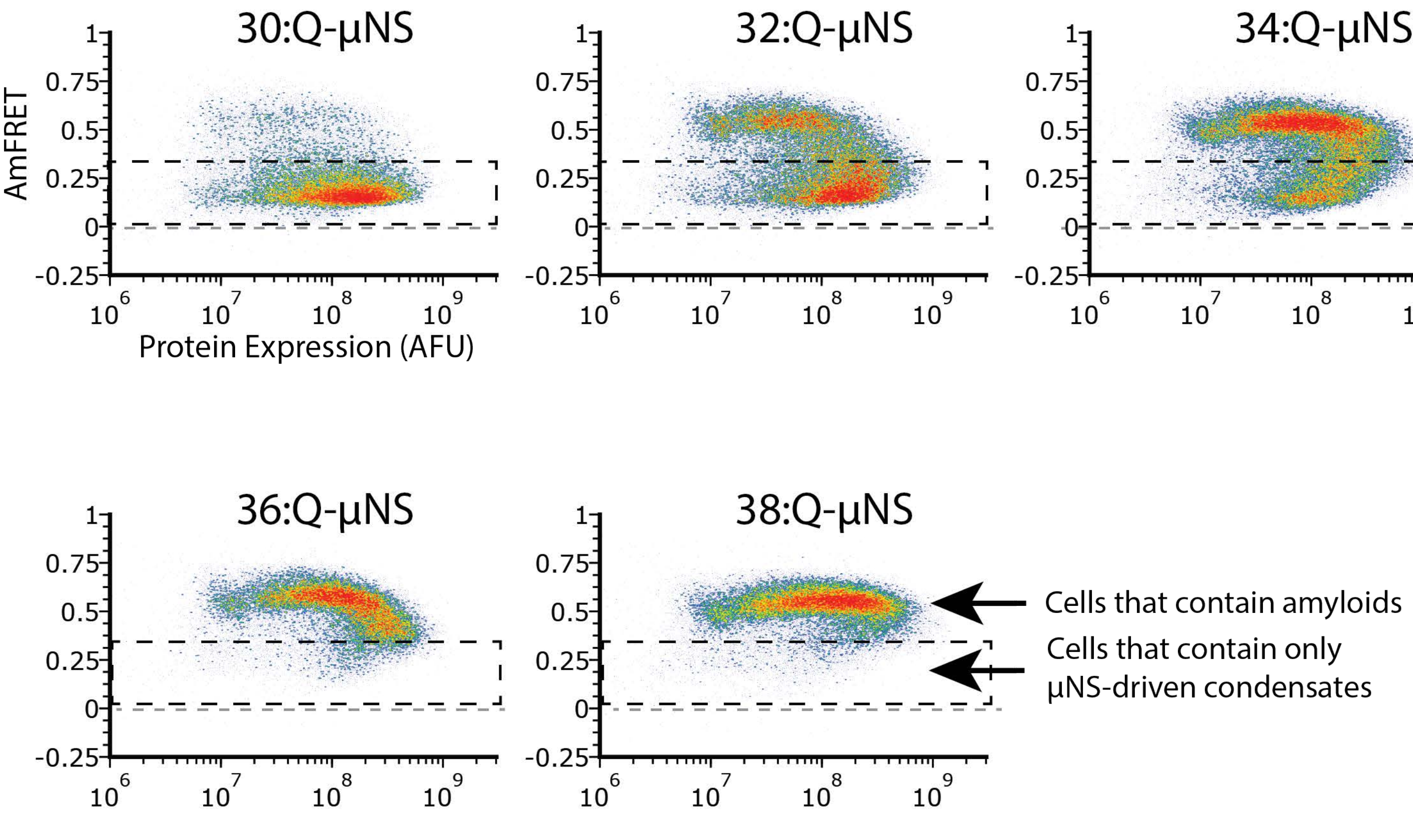

30:Q-μNS
32:Q-μNS
34:Q-μNS
36:Q-μNS
38:Q-μNS
AmFRET
1
0.75
0.5
0.25
0
-0.25
10^6
10^7
10^8
10^9
Protein Expression (AFU)
Cells that contain amyloids
Cells that contain only μNS-driven condensates

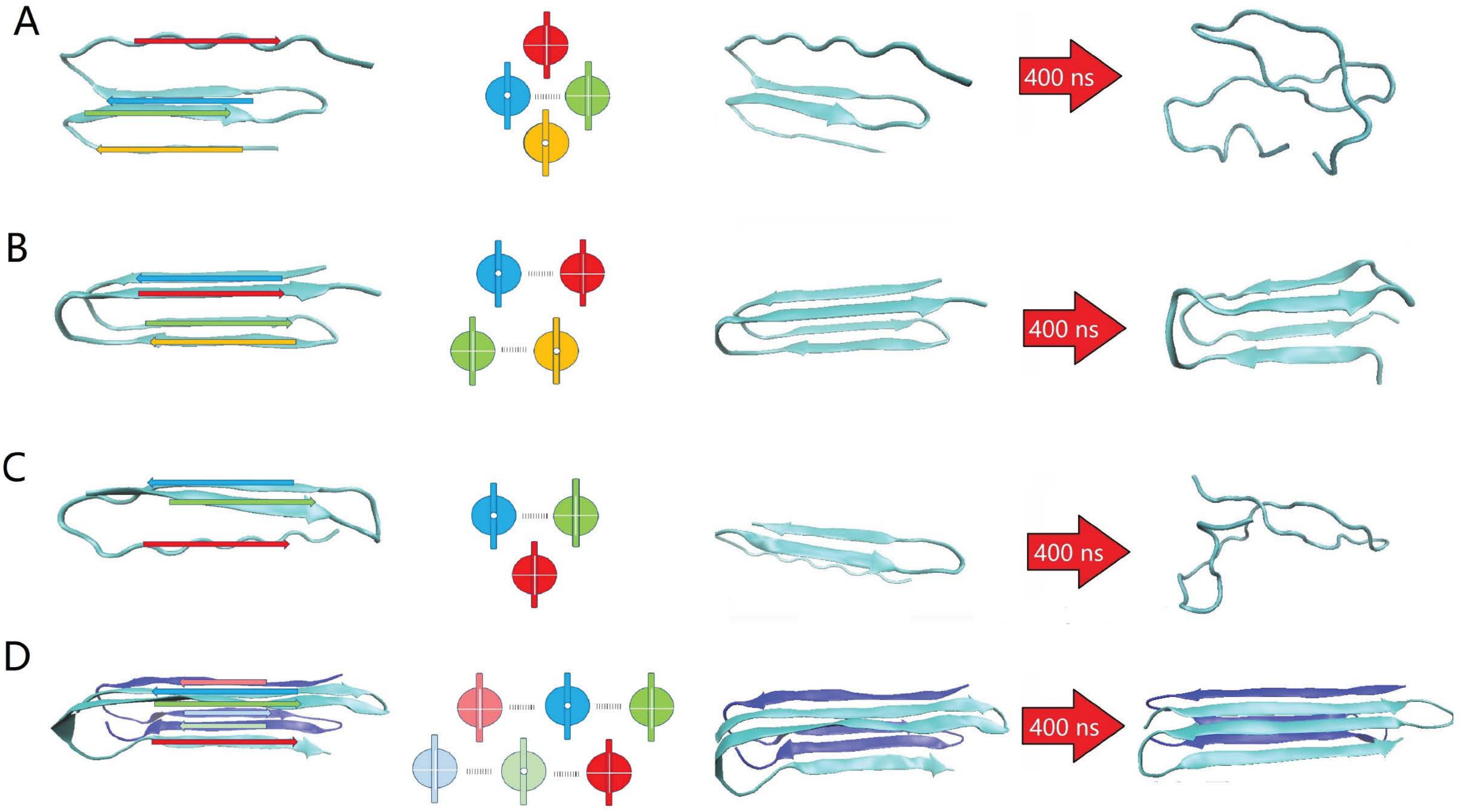
A
400 ns
B
400 ns
C
400 ns
D
400 ns
