## Supplementary material for "The polyglutamine amyloid nucleus in living cells is a monomer with competing dimensions of order": Table S2

*Table S2. Summary of results from publically available APR and aggregation propensity predictor webservers. All webservers were available publicly as of April 2021, and run with default parameters. Note that no methods distinguish between the experimentally determined non-amyloidogenic protein $Q_4N(60)$ and the amyloidogenic proteins $Q_3N(60)$ and $Q_5N(60)$. When available, quantitative outputs have been provided.*

| | | Amino Acid Sequence | | | | | |
|---|---|---|---|---|---|---|---|
| Application Name | Link | $Q(60)$ | $N(60)$ | $Q_3N(60)$ | $Q_4N(60)$ | $Q_5N(60)$ | Reference |
| AGGRESCAN[a] | http://bioinf.uab.es/aggrescan | non-amyloid (-1.231) | non-amyloid (-1.302) | non-amyloid (-1.249) | non-amyloid (-1.245) | non-amyloid (-1.243) | Chonchillo-SolÃ© et al. (2007) |
| WALTZ | https://waltz.switchlab.org | non-amyloid | non-amyloid | non-amyloid | non-amyloid | non-amyloid | Maurer-Stroh et al. (2010) |
| FoldAmyloid[b] | http://bioinfo.protres.ru/fold-amyloid/ | non-amyloid (19.23) | non-amyloid (18.49) | non-amyloid(19.05) | non-amyloid(19.08) | non-amyloid(19.11) | Garbuzynskiy et al. (2010) |
| 3D Profile (ZipperDB)[c] | http://services.mbi.ucla.edu/zipperdb/ | non-amyloid (-38.715) | non-amyloid (-42.280) | non-amyloid (-30.184) | non-amyloid (-40.875) | non-amyloid (-41.11) | Thompson et al. (2006) |
| Amylogram[d] | http://smorfland.uni.wroc.pl/shiny/AmyloGram/ | non-amyloid (0.0672) | non-amyloid (0.0672) | non-amyloid (0.0672) | non-amyloid (0.0672) | non-amyloid (0.0672) | Burdukiewicz et al. (2017) |
| ANuPP[e] | https://web.iitm.ac.in/bioinfo2/ANuPP/ | non-amyloid (19.00) | non-amyloid (15.69) | non-amyloid (13.18) | non-amyloid (14.18) | non-amyloid (14.43) | Prabakaran et al. (2020) |
| TANGO[f] | http://tango.crg.es/ | non-amyloid (0) | non-amyloid (0) | non-amyloid (0) | non-amyloid (0) | non-amyloid (0) | Fernandez-Escamilla et al. (2004) |
| SecStr | http://biophysics.biol.uoa.gr/SecStr/ | non-amyloid | non-amyloid | non-amyloid | non-amyloid | non-amyloid | Hamodrakas et al. (2007) |
| BETASCAN | http://betascan.csail.mit.edu/ | non-amyloid | non-amyloid | non-amyloid | non-amyloid | non-amyloid | Bryan et al. (2009) |
| Net-CSSP*[g] | http://cssp2.sookmyung.ac.kr/index.html | non-amyloid (0.068) | non-amyloid (0.021) | non-amyloid (0.066) | non-amyloid (0.063) | non-amyloid (0.067) | Kim et al. (2009) |
| AmyloidMutants | http://amyloid.csail.mit.edu/ | amyloid | non-amyloid | non-amyloid | non-amyloid | non-amyloid | O'Donnell et al. (2011) |
| PASTA 2[h] | http://old.protein.bio.unipd.it/pasta2/ | non-amyloid (1.095657) | non-amyloid (0.087910) | non-amyloid (1.448978) | non-amyloid (1.095657) | non-amyloid (1.095657) | Walsh et al. (2014) |
| GAP[i] | http://www.iitm.ac.in/bioinfo/GAP/ | amyloid (1.000) | amyloid (0.998) | amyloid(0.99925) | amyloid (0.9994) | amyloid (0.99956167) | Thangakani et al. (2014) |
| MetAmyl[j] | http://metamyl.genouest.org/ | amyloid (0.556610) | non-amyloid (0.493100) | amyloid (0.554175) | amyloid (0.552000) | amyloid (0.554100) | Emily et al. (2013) |
| Budapest Amyloid Predictor | https://pitgroup.org/bap/ | non-amyloid | non-amyloid | non-amyloid | non-amyloid | non-amyloid | Keresztes et al. (2020) |
| iAMY-SCM[k] | http://camt.pythonanywhere.com/iAMY-SCM | amyloid (697.00) | amyloid (550.00) | amyloid (490.59) | amyloid (533.36) | amyloid (561.86) | Charoenkwan et al. (2020) |

* Method correctly identified sequence $Q_4N(60)$ as having a lower amyloid propensity than $Q_3N(60)$ or $Q_5N(60)$.

[a] AGGRESCAN uses a fixed table of values to calculate the amino-acid propensity value ($a^3v$). If the value of $a^3v$ reaches the Hot Spot Threshold (-0.02) the sequence is predicted to be an aggregation prone protein.

[b] FoldAmyloid was run with default profile-value of 21.4, with a scale of 8Å for expected number of contacts, and an averaging frame of 5 residues.

[c]3D Profile (ZipperDB) uses sequential k-mers, with an energetic threshold of -23 kcal/mol or lower contributing to amyloid potential.

[d]AmyloGram results are in the form of a probability of an amyloid forming.

[e]ANuPP provides unitless scores.

[f]TANGO default parameters are no protection on the terminii, pH of 7, temperature of 298.15 Kelvin, ionic strength of 0.02M, and a concentration of 1M. The Agg parameter is the main result and gives the tendency for beta-sheet aggregation.

[g]CSSP2 results used the single network profile to calculate beta-sheet propensity.

[h]PASTA2 uses default values of 20 for Top Pairing Energy, and energy threshold of -5 (1.0 energy unit is equivalent and 1.192 Kcal/mol) using the Regular Algorithm (self aggregating) with a TPR (sensitivity) of 43.6% and a FPR (1-specificity) of 5.1% parameters. Best energy reported.

[i]Generalized Aggregation Proneness (GAP) calculates the amyloid probability for each hexamer in the full-length protein. Mean amyloid probability for each protein is reported.

[j]MetAmyl Prediction of Amyloid fragments uses a logistic regression model; the final score of each 6-residue sequence is the probability that k-mer will form an amyloid fiber. Note that while none of the hexapeptides reached the 0.6 probability threshold, the detailed profile view identified all of them as 'X - Amylogenic amino acid' even when the final probability was below 0.5, as in the case of sequence $N(60)$.

[k]Summary of improved Prediction and Analysis of Amyloid Proteins Using Scoring Card Method (iAMY-SCM) uses a default unitless threshold of 288.5625 to predict whether a given protein sequence is an amyloid protein.
